# Texture-like representation of objects in human visual cortex

**DOI:** 10.1101/2022.01.04.474849

**Authors:** Akshay V. Jagadeesh, Justin L. Gardner

## Abstract

The human visual ability to recognize objects and scenes is widely thought to rely on representations in category-selective regions of visual cortex. These representations could support object vision by specifically representing objects, or, more simply, by representing complex visual features regardless of the particular spatial arrangement needed to constitute real world objects. That is, by representing visual textures. To discriminate between these hypotheses, we leveraged an image synthesis approach that, unlike previous methods, provides independent control over the complexity and spatial arrangement of visual features. We found that human observers could easily detect a natural object among synthetic images with similar complex features that were spatially scrambled. However, observer models built from BOLD responses from category-selective regions, as well as a model of macaque inferotemporal cortex and Imagenet-trained deep convolutional neural networks, were all unable to identify the real object. This inability was not due to a lack of signal-to-noise, as all of these observer models could predict human performance in image categorization tasks. How then might these texture-like representations in category-selective regions support object perception? An image-specific readout from category-selective cortex yielded a representation that was more selective for natural feature arrangement, showing that the information necessary for object discrimination is available. Thus, our results suggest that the role of human category-selective visual cortex is not to explicitly encode objects but rather to provide a basis set of texture-like features that can be infinitely reconfigured to flexibly learn and identify new object categories.

**Significance Statement:** Virtually indistinguishable metamers of visual textures, such as wood grain, can be synthesized by matching complex features regardless of their spatial arrangement (1–3). However, humans are not fooled by such synthetic images of scrambled objects. Thus, category-selective regions of human visual cortex might be expected to exhibit representational geometry preferentially sensitive to natural objects. Contrarily, we demonstrate that observer models based on category-selective regions, models of macaque inferotemporal cortex and Imagenet-trained deep convolutional neural networks do not preferentially represent natural images, even while they are able to discriminate image categories. This suggests the need to reconceptualize the role of category-selective cortex as representing a basis set of complex texture-like features, useful for a myriad of visual behaviors.

## Introduction

Images engineered to have the same complex visual features as natural images can appear metameric (4, 5), i.e. perceptually indistinguishable though physically different from the original natural image. In particular, by matching image statistics (1, 2), including the pairwise correlations of orientation and spatial frequency filters (6) or the pairwise inner products of feature maps from Imagenet-trained deep convolutional neural networks (dCNNs) (7–9), it is possible to synthesize images which appear indistinguishable from the corresponding natural image, despite having a spatially scrambled arrangement of features (3, 10). This metamer synthesis approach is particularly effective for visual textures, such as bark, gravel, or moss, which contain complex visual features that are largely homogeneous over space (11–15). The synthesis approach has been used to study the phenomenon of crowding (16) in peripheral vision (10, 17, 18), where observers can fail to bind visual features to corresponding objects when presented in visual clutter (19, 20). The neural representation of complex visual features has also been studied with texture synthesis approaches (21–25).

However, images which contain inhomogeneous visual features, such as those of objects or natural scenes, are perceptually distinct from synthesized images which contain the same complex visual features but in scrambled spatial arrangements (14, 26, 27). This suggests that the underlying neural representation of objects and natural scenes, in contrast to that of textures, is sensitive to the particular spatial arrangement of features found in objects and scenes in the natural world.

The lateral occipital and ventral visual cortex of humans is a potential cortical substrate for such representations which distinguish objects and natural scenes from synthesized, scrambled counterparts. Studies of early visual cortical representations suggest that sensitivity to the mid-level visual features contained in texture images can be found in areas V2 (21–23) and V4 (24, 25, 28), while category-selective representations in higher-level visual cortical areas within lateral occipital cortex (LO) and ventral temporal cortex (VTC) are informative for decoding object categories (29–35) and predicting object categorization behavior (30, 36, 37). Thus, one might hypothesize that cortical representations in category-selective regions underlie the perceptual ability to discriminate natural object images from synthesized images containing spatially scrambled arrangements of complex visual features (38–43).

However, evidence of texture bias in deep convolutional neural network models (44–46) of the ventral visual stream suggests that it is possible to encode features useful for discriminating visual object categories without explicitly encoding the global arrangement of those features. Imagenet-trained (47) deep convolutional neural networks (dCNNs) have achieved state-of-the-art performance at modeling ventral visual cortical representations (48–52), but unlike human perception, these dCNN models are biased towards texture rather than shape information (53–56). That is, when texture and shape information conflict, Imagenet-trained dCNNs frequently prioritize the texture information, whereas humans reliably prioritize shape. The evidence of texture-bias in dCNNs but not in human perception suggests two possible hypotheses: either dCNNs may not be an accurate model of primate ventral visual cortex, particularly with regards to the processing of shape information, or the representations in ventral visual cortex are also texture-like (57) and are therefore insufficient to account for humans’ perceptual ability to discriminate natural scenes from synthesized images containing the same complex visual features in scrambled arrangement.

One approach to address whether human ventral visual cortex explicitly represents the spatial arrangement of features that defines an object or natural scene is to examine the representational dissimilarity between natural images and synthesized images that have the same visual features but are scrambled. Prior studies employing this scrambling technique have been instrumental for the discovery of complex feature selectivity and invariance in high-level visual cortical areas (28, 57–61). However, these studies have often used methodology for scrambling visual features which confounds sensitivity to complex visual features with sensitivity to the spatial arrangement of those features. For example, Grill-Spector et al. (58) reported object sensitivity in LO by contrasting the BOLD response to natural object images with the response to grid-scrambled images, a scrambling approach which breaks up complex visual features. Similarly, Rust and DiCarlo (28) demonstrated enhanced selectivity and tolerance to objects in macaque inferotemporal (IT) cortex by comparing neural responses to natural object images with responses to synthesized scrambles containing only low and mid-level visual features. Thus, it remains undetermined whether cortical representations in category-selective regions of visual cortex are merely sensitive to the presence of complex visual features or whether they are also sensitive to the spatial arrangement of those features. Furthermore, prior research has primarily examined the magnitude of response averaged across entire cortical areas (21, 58–60), rather than the multivariate patterns of population activity that might provide much richer featural representations. Finally, this work has largely overlooked the link between these cortical representations and the perceptual behaviors which they support.

In the present study, we compared the ability of human observers, dCNN models, and cortical representations in category-selective regions of human visual cortex to discriminate natural images from synthesized images containing scrambled complex features. We sought to avoid the confound between feature complexity and spatial arrangement by adapting a hierarchical, spatially constrained image synthesis algorithm that allows control over the complexity of features and the spatial extent over which those features can be scrambled in the synthesized output (7, 8). We used these images as stimuli to assess both behavioral and cortical sensitivity. We found that human observers are highly sensitive both to the complexity of visual features as well as the spatial arrangement of those features in images of objects. In stark contrast, Imagenet-trained dCNN models were only sensitive to the complexity of features, not to the arrangement of those features, regardless of whether the natural image is of an object or texture. Similarly, we found that the cortical representations in category-selective regions of human visual cortex were also texture-like, i.e. non-selective for the natural arrangement of complex features in object images. Taken in sum, these results demonstrate that visual cortical representations and those of dCNNs, are insufficient to explain human sensitivity to natural arrangements of complex features in objects. This suggests that further stages of computation are required to account for human perceptual discriminability of natural from synthesized scrambled object images (62). While we found that no general linear readout of cortical response could better discriminate the natural image, image class specific linear readouts could be constructed which improved the discriminability of natural images on a image-by-image basis. This suggests that information about the natural spatial arrangement of features is available in the cortical representation and could be used by experience-dependent readout mechanisms of visual cortical representations.

## Results

### Human Perceptual Sensitivity for Natural vs Synthesized Images

To assess perceptual sensitivity to the complexity and spatial arrangement of features, we used an oddity detection task, in which subjects were presented with 3 images on each trial (1 natural, 2 synthesized) and asked to choose the odd-one-out, i.e. the image which appeared the most different from the others. Images subtended 8 degrees in diameter and were centered 6 degrees from the center of fixation. On each trial, both synthesized images (“synths”) were matched to the features of the natural image at a particular level of feature complexity and spatial arrangement constraint. To synthesize images, we extracted feature maps from various VGG-19 layers (e.g. pool1, pool2, pool4) in response to a natural image, then computed the pairwise dot products between each pair of feature maps (Gramian) within subregions of different levels of spatial constraint (from least spatially constrained, 1×1, where features could be spatially scrambled across the whole image to most spatially constrained, 4×4, in which features could only be scrambled within 16 subregions of the image) (**Fig. 1A**). Then, we iteratively optimized the pixels of a randomly initialized white noise image to minimize the mean squared error between its Gramian feature representation and that of the original. We will refer to the feature complexity of a synth to indicate the latest dCNN layer whose features were extracted and matched to the original. We will use the term spatial constraint or spatial arrangement to indicate the size of the spatial pooling regions within which those features were matched. Across trials, we varied the feature complexity of the synths as well as the degree to which the spatial arrangement of those features was constrained (**Fig. 1B**). We will use the term image class to refer to the set of images including a given natural image and all its feature-matched synths.

**Figure 1.**
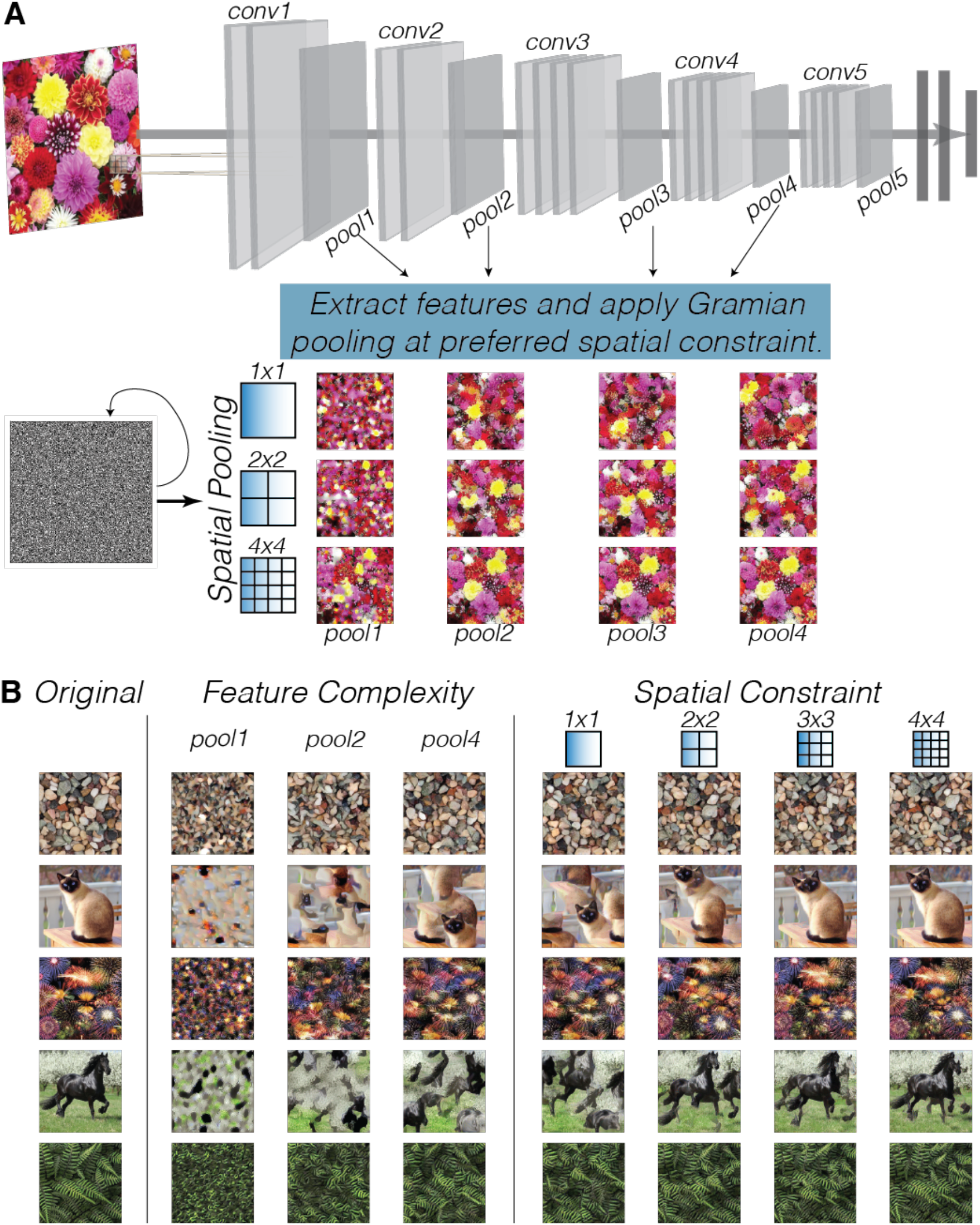
Image synthesis algorithm and example synths. (A) Schematic of deep image synthesis algorithm. We pass a natural image into Imagenet-trained VGG19 model and extract intermediate layer activations from layers pool1, pool2, and/or pool4. Then, we compute the Gramian constrained within spatial pooling regions (1×1, 2×2, 3×3, or 4×4). To synthesize, we iteratively update pixels of a random seed image using gradient descent to match spatially pooled Gramian of original image. (B) Example natural images and feature-matched synths, varying in feature complexity (columns 2-4, fixed at 1×1 spatial constraint), and varying in spatial constraint (columns 5-8, fixed at pool4 complexity).

We found that human observers were less able to detect the natural image among synths with more complex features. To determine the effect of feature complexity on the discriminability of natural and synthesized images, we analyzed oddity detection performance as a function of the feature complexity of the synths, pooled across all observers and averaged across all spatial constraints. We reasoned that this would be informative of which features are utilized in the perception of natural images of objects. We found that increasing the feature complexity of the synths resulted in a significant decline *(Linear mixed effects model: b=-0.103, SE=0.007, p<0.001, 95% CI=[−0.116, −0.090], N=87)* in the proportion of trials where the natural image was chosen as the oddity (**Fig. 2C**, note downward slope of purple line), suggesting that human observers are perceptually sensitive to the complexity of visual features in object images.

**Figure 2.**
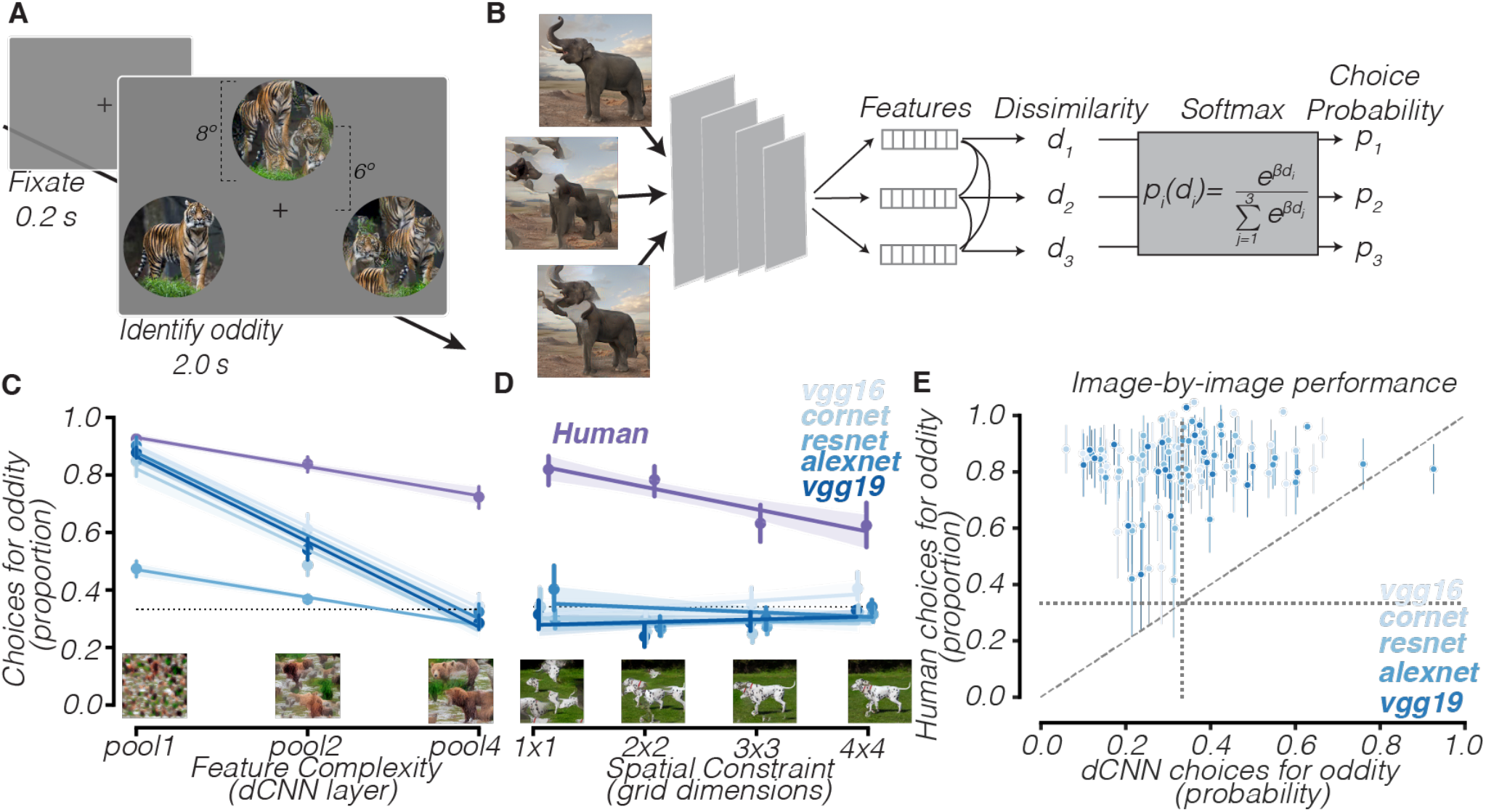
Human perception of objects is sensitive to feature complexity and spatial arrangement; dCNNs are insensitive to spatial arrangement. (A) Oddity detection task. Subjects see 3 images, one natural and two synths, and choose the odd-one-out. (B) Schematic of dCNN observer model fit to oddity detection task. (C) dCNN performance (blue) compared to human performance (purple) as a function of synths’ feature complexity. Horizontal dotted line represents chance-level. Example synths from each feature complexity level are shown at the bottom. (D) dCNN performance (blue) compared to human performance (purple) as a function of synths’ spatial constraints. Horizontal dotted line represents chance-level. Example synths from each spatial constraint level are shown at the bottom. (D) dCNN performance vs human performance, class-by-class, for 1×1 pool4 condition. Diagonal dashed line is line of equality. Vertical/horizontal dotted lines represent chance-level.

We found that human observers were also less able to detect the natural image among synths with more constrained feature arrangement. To assess perceptual sensitivity for the spatial arrangement of visual features, we analyzed oddity detection performance as a function of the spatial constraints in the synths, pooled across all observers, fixing feature complexity at the highest level (pool4). We reasoned that if observers were no less able to detect the natural image among synths whose features were constrained within small subregions of the image (e.g. 4×4 condition) as compared to synths whose features were scrambled across the entirety of the image (1×1 condition), this would demonstrate that humans are insensitive to the arrangement of complex visual features. We found instead that the proportion of trials where subjects selected the natural image as the odd-one-out significantly decreased *(Linear mixed effects model: b=-0.087, SE=0.011, p<0.001, 95% CI=[−0.108, −0.066])* as the arrangement of object features was more strongly constrained (**Fig. 2D**, note downward slope of purple line). This pattern of behavior was consistent regardless of whether behavioral data were collected in-lab with fixation enforced (**Fig. S3**) or online. These findings suggest that human observers’ perception of objects is not only sensitive to the presence of complex visual features but also the spatial arrangement of those features.

### dCNN Observer Models

To compare the behavior of dCNN models to that of human observers, we constructed an observer model that uses dCNN features to perform the oddity detection task (**Fig. 2B**). On each trial, we first extracted a feature vector corresponding to each image, from the last convolutional layer of an Imagenet-trained dCNN. We then computed the Pearson distance between each pair of feature vectors. To determine which image is most different from the other two, we computed the dissimilarity of each image by averaging the distance from each image to each of the two other images. These dissimilarities were then transformed into choice probabilities using a softmax function, with one free parameter controlling how sensitive the model is to dissimilarity differences, which was estimated to maximize the likelihood of human observers’ choices. We evaluated the performance of five different Imagenet-trained dCNNs: VGG-19 (63) (same model used for image synthesis), CORnet-Z (64), VGG-16 (63), ResNet-18 (65), and AlexNet (46). We hypothesized that if dCNNs are an accurate model of human object perception, they ought to match human performance at this oddity detection task, in terms of sensitivity to both feature complexity and spatial arrangement.

When synths contained more complex features, we found that dCNN observer models were less able to identify the natural image as the odd-one-out. We compared each model’s performance to human oddity detection performance as a function of the feature complexity of the synthesized images contained, averaged across all spatial constraints. Like human observers, almost all models were very likely to select the natural image when presented among synths containing only low-level visual features (**Fig. 2C,** pool1). However, increasing the feature complexity of the synths resulted in a steep decline in the probability of selecting the natural image *(Linear mixed effects model: b=-0.249, SE=0.001, p<0.001, 95% CI=[−0.251, −0.247]),* such that all dCNN observer models were not significantly more likely than chance *(b=-0.038, SE=0.012, p=0.998, 95% CI=[−0.061, −0.014])* to identify the natural object image among two synths containing complex visual features (**Fig. 2C**, pool4). This contrasts with human observers, whose frequency of selecting the natural image as the oddity did decline with increasing feature complexity but was still significantly above chance even at the highest level of feature complexity (**Fig. 2C**, note purple line), and suggests that dCNNs are sensitive only to the presence of complex visual features and not to their arrangement.

When synths contained complex visual features, regardless of how scrambled those features were, we found that dCNN observer models were unable to detect the natural image. To determine whether dCNN observer models are sensitive to the spatial arrangement of features or merely to the presence of the complex features which make up objects, we analyzed oddity detection performance as a function of spatial pooling region size, for the complex (pool4) feature condition alone to isolate the effect of spatial arrangement when fixing feature complexity at the highest possible level (**Fig. 2D**). Whereas human observers detected the natural image most frequently when feature arrangement was least constrained (**Fig. 2D,** 1×1, note purple line) and less frequently as feature arrangement became more constrained, all five dCNN observer models were not significantly more likely than chance to select the natural image in any condition *(b=-0.038, SE=0.012, p=0.998, 95% CI=[−0.061, −0.014])* and were insensitive to variation in spatial arrangement *(b=0.003, SE=0.001, p=0.02, 95% CI=[0.000, 0.005])* (**Fig. 2D**, note flat blue lines). Thus, unlike human observers, who reliably report that the natural image stands out among synthesized images, dCNN models are unable to identify the natural image as the odd-one-out even when presented among images whose features are scrambled across the entire image (1×1). Across a large sample of object image classes (**Fig. 2E**), we found that humans were significantly more likely than all Imagenet-trained dCNN models we tested to choose the natural image as the oddity when presented among synths containing scrambled (1×1) complex (pool4) visual features *(VGG16: t=12.84, p<0.001; VGG19: t=14.93, p<0.001; CORnet: t=14.04, p<0.001; ResNet18: t=17.04, p<0.001; AlexNet: t=9.47, p<0.001).* Thus, given that dCNNs were at floor performance regardless of how well-constrained the arrangement of features was, we hypothesized that what distinguishes humans’ perception of objects from that of dCNNs is not the ability to detect complex visual features, but rather a selectivity for the particular spatial arrangement of features found in natural objects.

### Category-level oddity detection task

These behavioral and modeling results demonstrate that dCNN features are insensitive to the arrangement of complex features, suggesting that dCNNs do not represent objects but instead contain a texture-like representation of disjointed complex visual features. One implication of this is that object categorization, the task that Imagenet-trained dCNNs are optimized to perform, does not require an explicit representation of objects, merely a representation of the complex features that make up objects. To test this, we compared the performance of human observers and dCNN observer models in a category-level oddity detection task (**Fig 3A**). On each trial, observers saw three different natural images - two of which belonged to the same category, while the third image contained an image from a different category - and were instructed to choose the odd-one-out. To make this task comparable to the previous task, subjects were instructed to select the image which appeared most different from the other two and were not explicitly directed to choose the image which belonged to a different category. We hypothesized that human observers’ behavior on this category-level oddity detection task would be predicted well by the performance of dCNN observer models.

**Figure 3.**
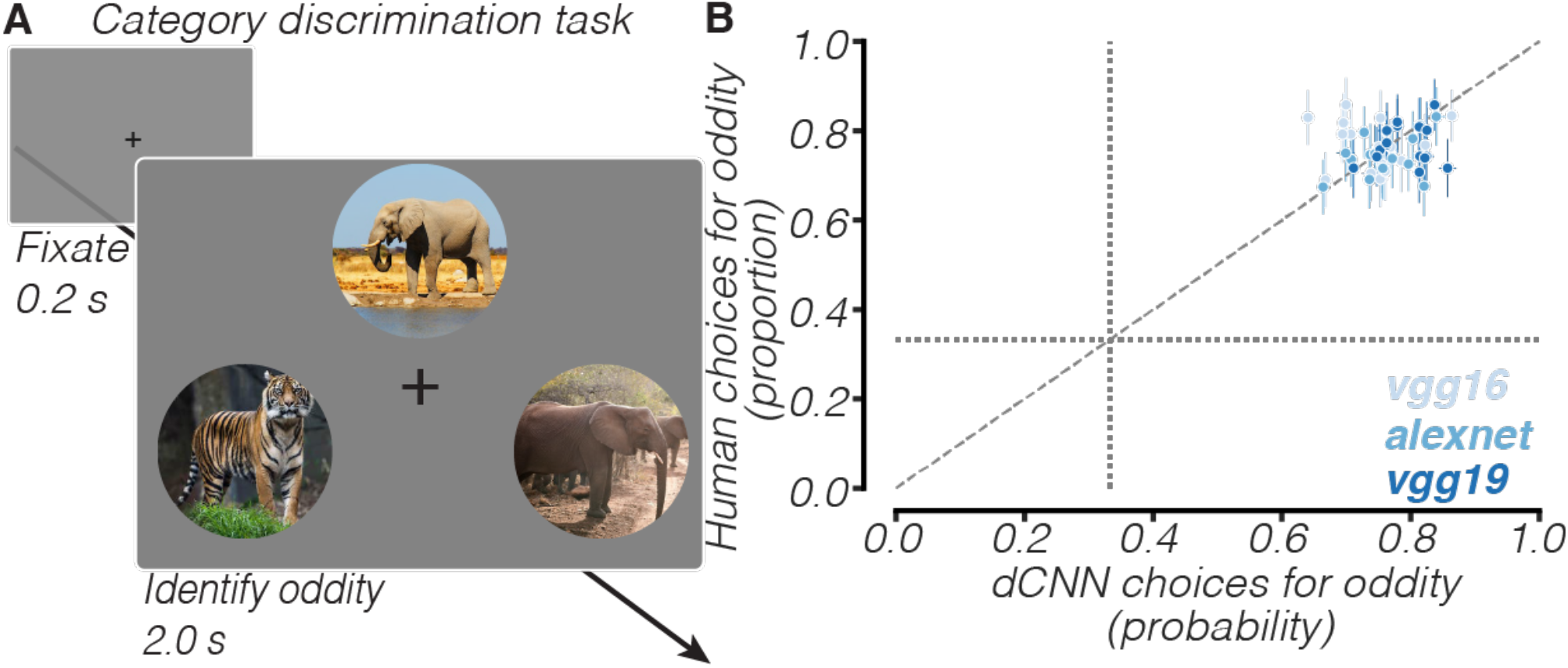
dCNN observer models match human performance in a category-level oddity task. (A) Category-level oddity detection task. On each trial, subjects see 3 natural images – 2 of the same category, 1 from a different category – and choose the odd-one-out. (B) dCNN vs human performance, category-by-category. Diagonal dashed line is line of equality. Vertical/horizontal dotted lines represent chance-level.

Indeed, we found that human observers’ behavioral performance at category-level oddity detection was matched by the performance of dCNN observer models. We compared the frequency with which dCNNs selected the image belonging to the odd-category-out to that of human observers. We found that across a variety of object categories, there was no significant difference *(t=1.16, p=0.26, n=16)* in performance between human observers and dCNN observers (**Fig 3B**). This suggests that dCNN representations, although insensitive to feature arrangement, are informative for discriminating categories that differ in their visual features.

### Human visual cortex

To assess the ability of the human visual cortex to discriminate between natural and synthesized object images, we measured BOLD responses in the brains of 7 human observers, while subjects passively viewed natural and synthesized (1×1, pool4) images from 10 different image classes. Images were presented for 4 seconds, subtending 12 degrees, centered 7 degrees to the left and right of fixation. We estimated trial-averaged responses to each individual image using a generalized linear model (66, 67). We analyzed data from 13 visual cortical areas, including V1, V2, V3, and hV4, which were retinotopically defined (68), mid-fusiform (mFus), posterior fusiform (pFus), inferior occipital gyrus (IOG), transverse occipital sulcus (TOS), and collateral sulcus (CoS), which were functionally defined using a functional localizer (69), and lateral occipital cortex (LO), ventral visual cortex (VVC), posterior inferotemporal cortex (PIT), and ventromedial visual area (VMV), which were anatomically defined (70). We use the term early visual cortex to refer to V1, V2, V3, and hV4 and the terms category-selective regions or category-selective cortex to refer to mFus, pFus, IOG, TOS, CoS, VVC, PIT, LO, and VMV.

We were able to measure reliable patterns of BOLD activity in these 13 cortical areas in every subject. We estimated the split-half reliability of each voxel as the correlation between its responses, estimated on two different halves of image presentations. In each visual area, we selected the 100 voxels with the highest split-half reliability, to ensure our findings reflected a reliable signal rather than just noise. The mean reliability across all subjects of the 100 selected voxels in each of these visual areas exceeded the chance level derived from permutation testing in all analyzed visual areas (*R_V1_ = 0.70; R_V2_ = 0.68; R_V3_ = 0.63; RhV4 = 0.56; R_mFus_ = 0.30; R_pFus_ = 0.47; R_IOG_ = 0.42; R_CoS_ = 0.45; R_TOS_ = 0.44; R_LO_ = 0.58; R_VVC_ = 0.33; R_PIT_ = 0.33; R_VMV_ = 0.44).*

To determine whether visual cortical responses could support behavior in the oddity detection task, we constructed an observer model that used BOLD responses to choose the image which was most different (**Fig 4A**). On each trial, we first extracted a vector of voxel responses from a given visual area to each of the three images that the human saw, then computed the Pearson distance between each response vector, averaged together each pair of distances to estimate the representational dissimilarity of each image (mean distance from other two images), and then transformed the dissimilarities into choice probabilities using a softmax function. We evaluated the ability of this cortical observer model to identify the odd-image-out in both the category-level oddity detection task, where two objects of the same category were presented alongside a third object of a different category, and the natural-vs-synth oddity detection task, where two synths containing scrambled complex visual features were presented alongside a natural image.

**Figure 4.**
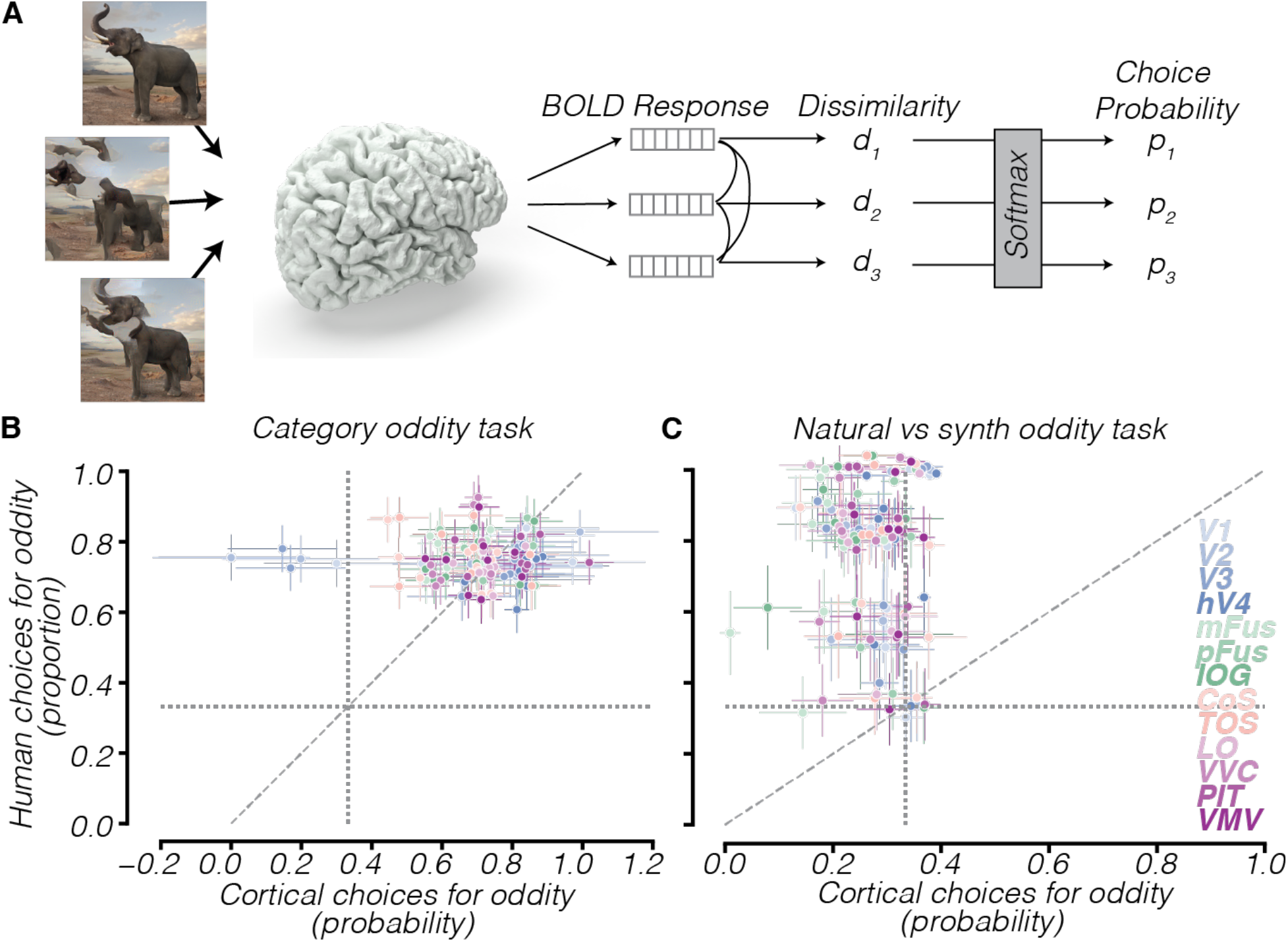
BOLD responses match human performance at category-level discrimination but not natural-vs-synth discrimination. (A) Schematic of cortical observer model which uses BOLD responses to perform oddity detection task. (B) Cortical observer model vs human performance on category-level oddity detection task, category-by-category. (C) Cortical observer models vs human performance, class-by-class, on natural-synth discrimination task. Diagonal dashed line is line of equality. Vertical/horizontal dotted lines represent chance-level.

In a category-level oddity detection task, human observers’ behavioral performance was matched by that of BOLD responses in category-selective regions of human visual cortex. Comparing the performance of the cortical observer model which makes use of BOLD responses from visual cortical regions to estimate oddity detection task performance to that of human observers, we found that there was no significant difference (*V1: t=-1.49, p=0.17; V2: t=-0.75, p=0.47; V3: t=-1.04, p=0.32; hV4: t=4.47, p=0.02; mFus: t=-4.05, p=0.03; pFus: t=-0.88, p=0.40; IOG: t=-1.18, p=0.27; CoS: t=-4.31, p=0.02; TOS: t=-1.46, p=0.18; LO: t=-1.94, p=0.09; VVC: t=-1.17, p=0.29; PIT: t=0.74, p=0.48; VMV: t=-1.63, p=0.14)* in the likelihood of selecting the odd-category-out between human behavior and human cortical responses in all visual cortical regions analyzed except three (hV4, mFus, and CoS, all marginally significant) (**Fig. 4B**). Early visual cortical regions were able to discriminate some categories, likely due to basic featural differences between categories, but not all categories (**Fig. 4B,** note blue points in top left). These results suggest that the BOLD responses we measured in human visual cortex contain useful information for discriminating between different categories.

When discriminating natural images from feature-matched scrambled synths, cortical responses were unable to match the performance of human observers. Across several different image classes, we found that all observer models constructed using responses from each visual area were significantly less likely (*V1: t=-5.89, p<0.001; V2: t=-6.25, p<0.001; V3: t=-6.21, p<0.001; hV4: t=-5.76, p<0.001; mFus: t=-8.70, p<0.001; pFus: t=-6.78, p<0.001; IOG: t=-6.36, p<0.001; CoS: t=-5.32, p<0.001; TOS: t=-6.66, p<0.001; LO: t=-5.93, p<0.001; VVC: t=-7.76, p<0.001; PIT: t=-5.24, p<0.001; VMV: t=-6.53, p<0.001)* to identify the natural object image compared to human observers (**Fig 4C**). This suggests that visual cortical responses, across distinct functional and anatomical areas including early visual cortex (V1, V2, V3, hV4), ventral temporal cortex (mFus, pFus, CoS, VMV, VVC, PIT) and lateral occipital cortex (LO, IOG, TOS) do not preferentially represent the natural arrangement of object features relative to scrambled arrangements containing the same complex visual features, which suggests that representations in category-selective regions of human visual cortex lack selectivity for natural feature arrangement.

### Model of Macaque Inferotemporal Cortex

To examine the ability of neurons in macaque inferotemporal (IT) cortex to perform this oddity detection task, in the absence of electrophysiological recordings in response to our synths, we evaluated a dCNN-based model of IT neuronal response. This model was constructed by linearly transforming the feature space from a late convolutional layer of an Imagenet-trained dCNN to maximize predictivity of a population of 168 IT sites (71), an approach which has yielded state-of-the-art performance at predicting out-of-sample IT neural responses (48, 72). This IT model explains on average across sites 51.8% of the variance in neural response to held-out images (72). We used the responses of these model IT neurons as inputs to an observer model constructed to perform the oddity detection task (**Fig 5A**). On each trial, the observer model computed the response of the model IT neurons to each of the 3 images, then computed the Pearson distance between each of these 3 response vectors, averaged each pair of distances together to compute the dissimilarity of each image, and then used a Softmax function to determine the probability of choosing each image. We evaluated two different dCNN models fit to IT response, one from the final convolutional layer of Alexnet and the other from VGG19 layer pool5.

**Figure 5.**
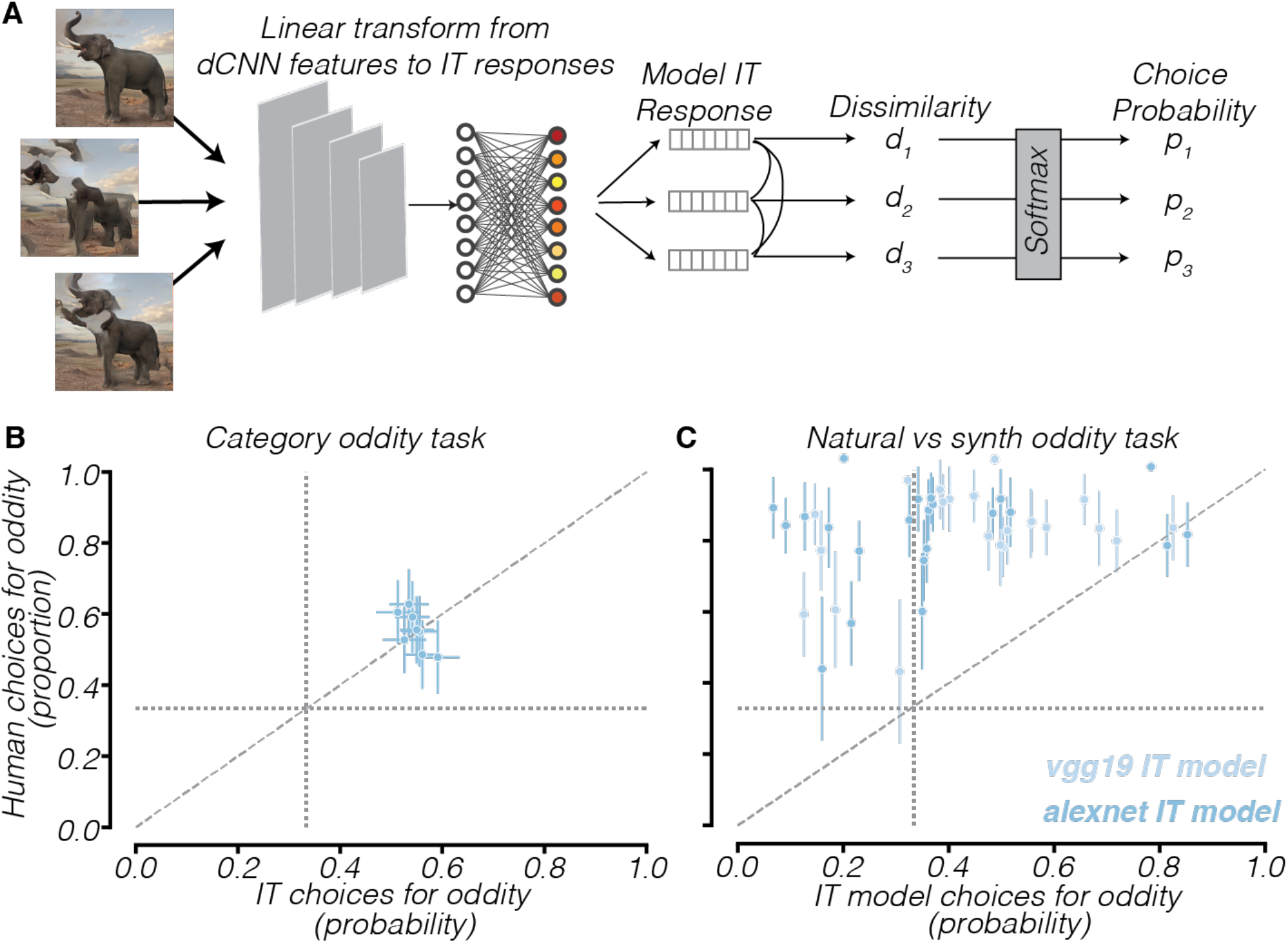
IT model matches human performance at category-level discrimination but not natural-vs-synth discrimination. (A) Schematic of IT observer model which uses model IT responses to perform the oddity detection task. (B) IT observer model vs human performance on category-level oddity detection task, category-by-category. (C) IT observer model vs human performance on natural-synth discrimination task, class-by-class. Diagonal dashed line is line of equality. Vertical/horizontal dotted lines represent chance-level.

In a category-level oddity detection task, macaque IT responses matched human performance. We used a dataset of macaque IT multiunit electrode recordings (71) to estimate category-level oddity detection performance. We presented human observers with the same images that were seen individually by the macaques. Then, we compared human performance at this oddity detection task to the estimated performance of a population of IT neural sites at performing this task and found that they were not significantly different (*t=-0.235, p=0.821)* (**Fig 5B**), suggesting that macaque IT contains feature representations useful for discriminating between images of different categories.

When discriminating natural images from synths with complex scrambled features, model IT neurons were far less likely than humans to choose the natural image as the oddity. We compared the oddity detection performance of this population of 168 IT model neurons to human behavioral performance, specifically for the condition in which subjects saw one natural image and two synths with complex (pool4) features in a scrambled (1×1) arrangement. We found that across a wide variety of image classes, humans were significantly more likely (*t=9.46, p<0.001, N=87)* than model IT neurons to pick the natural image (**Fig 5C**). This finding suggests that the model IT population does not differentially represent the natural arrangement of features compared to scrambled arrangements containing the same complex features.

### Representational geometry analysis

The discrepancy in behavior on the natural-vs-synth oddity detection task between human observers and dCNN models, category-selective regions of human visual cortex, and macaque IT models suggests a misalignment in the underlying representational geometries which give rise to the observed behaviors. Therefore, we directly analyzed the representational spaces of dCNNs, category-selective visual areas, and model IT, and compared them to the perceptual representational space, which we inferred from behavioral responses in an independent perceptual task.

Using a similarity judgment task, we found that the representations that underlie human object perception must be selective for the natural arrangement of features. To assay the representational geometry of humans’ perception of objects, we conducted an independent behavioral experiment in which we presented two pairs of images on each trial and asked observers to choose which pair was more internally dissimilar (**Fig. 6A**) (73, 74). With the responses from this experiment pooled across all observers, we used a modified version of Maximum Likelihood Difference Scaling (MLDS) (73) to estimate the perceptual distances between pairs of images. Whereas the original MLDS method estimates the position of different stimuli along a single dimensional axis to maximize the likelihood of the psychophysical responses (N free parameters for N stimuli), we directly estimated pairwise distances between each pair of stimuli (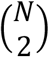 free parameters for N stimuli) to eliminate any assumptions about the dimensionality of the representational space. We visualized those distances by plotting the three images presented on a given trial of the oddity detection task (for category-level task, one image from one category and two images from other category; for natural-vs-synth task, one natural image and two synths containing complex (pool4) features in scrambled (1×1) arrangements) in a triangle, such that the length of the edges corresponded to the pairwise perceptual distances, a visualization which we will refer to as a triangular distance plot (**Fig 6**). We similarly visualized the representational geometry of the feature space represented by the final convolutional layer of each dCNN observer model as well as the representational geometry of human category-selective visual cortical regions and the representational geometry of a macaque IT model.

**Figure 6.**
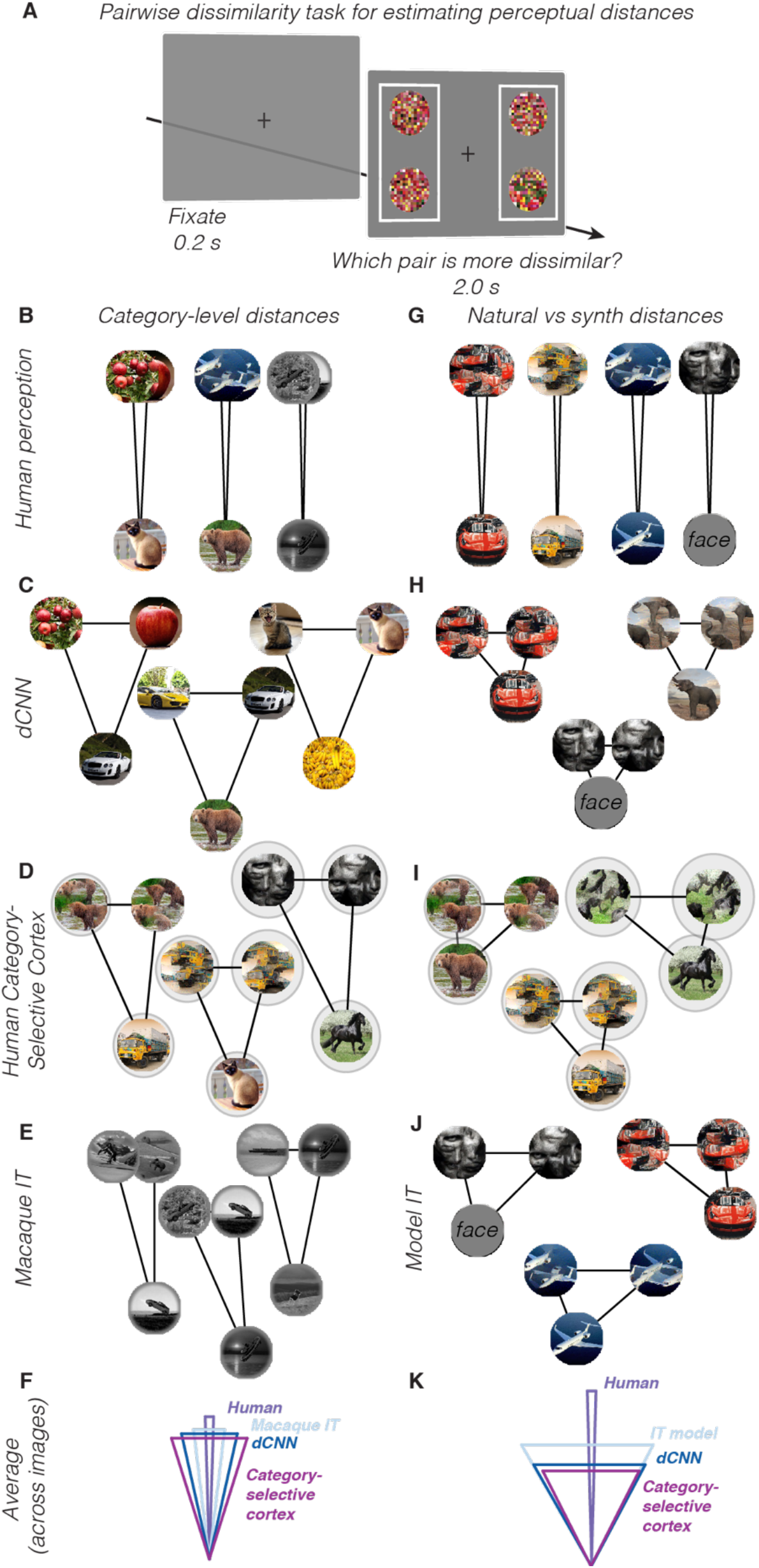
Representational geometry misalignment between humans and model/cortical representations for natural vs synth discrimination. (A) Pairwise dissimilarity judgment task. Subjects saw two pairs of images and reported which pair was more dissimilar. Responses from this task were used to estimate the relative perceptual distances between pairs of images. (B) Perceptual distances, estimated via modified MLDS, between images of different categories and images of the same category, for 3 example categories. (C) dCNN representational distances between images of different categories and images of the same category, for 3 example categories. (D) Cortical representational distances between images of different categories relative to images of same category, for 3 example categories. (E) Macaque IT representational distances between images of different categories and images of same category, for 3 example categories. (F) Triangular distances averaged across all categories. (G-K) Same as A-E but for natural-synth representational distance relative to synth-synth distances.

For the category-level discrimination, we found that the representational geometry of human visual perception was well aligned with that of dCNNs, human category-selective visual cortex, and macaque IT neurons. The estimated perceptual distance between two images of two different categories significantly exceeded the estimated perceptual distance between two images of the same category (*t=93.49, p<0.001)* (**Fig. 6B**, note narrow triangles). Similarly, for dCNN models (**Fig. 6C**), category-selective regions in human visual cortex (**Fig. 6D**), and macaque IT cortex (**Fig. 6E**, the representational distance between images of different categories significantly exceeded the representational distance between images of the same category. This alignment of representational geometry (**Fig. 6F**) explains why human performance on the category-level oddity detection task was well matched by that of dCNN observer models (**Fig. 3B**), category-selective regions of visual cortex (**Fig. 4B**), and macaque IT neurons (**Fig. 5B**).

However, for the natural-vs-synth discrimination, we found a misalignment between the representational geometry of human visual perception and those of dCNNs, category-selective regions of human visual cortex, and model IT. For human visual perception, we found that the estimated representational distance between the natural image and the synthesized images was significantly greater (*t=69.25, p<0.001)* than the representational distance between two different synths of the same class (**Fig. 6G**, note the narrow triangles), which reflects a selectivity for the natural intact arrangement of object features. These results suggest that the representations directly underlying object perception must be selective for the arrangement of object features.

In contrast, for all dCNN models we tested, we found that the representational distance between the natural image and the synthesized images was not significantly different (*t=-0.46, p=0.65, N=87)* from the representational distance between two different synthesized scrambled images (**Fig 6H**, note approximately equilateral triangles). This representational geometry analysis suggests that dCNN observer models are non-selective for natural arrangements of object features, in stark contrast to human observers who are highly selective for natural object arrangement (**Fig 6K**).

Similarly, representations in category-selective regions of human visual cortex and in models of macaque IT neurons were non-selective for the natural arrangement of objects, unlike perception. Using BOLD responses from category-selective regions, we found that the representational distance between a natural image and a feature-matched synth was not significantly greater than the representational distance between two different synths of the same image class (*t=-1.52, p=0.17)* (**Fig 6I**). This representational geometry of category-selective visual cortical regions differs greatly from that of human perception, which is highly selective for the natural arrangement of object features (**Fig 6K**). In models of macaque IT neurons, we also found that the representational geometry was non-selective for spatial arrangement. Although one might assume this analysis is bound to come to the same conclusion as the prior analysis of the unweighted dCNN representation, representational distances are not invariant to linear transformations, so linearly weighting dCNN features to fit model IT responses could have yielded a representation selective for feature arrangement. However, we found that for this model IT population, the representational distance between a natural image and a feature-matched synth was not significantly different (*t=0.78, p=0.44)* from the representational distance between two different synths (**Fig 6J**). These findings demonstrates that representations in category-selective regions of visual cortex and in a model of macaque IT cortex are non-selective for the natural arrangement of features and that the representational geometry of visual cortex is therefore misaligned with that of human visual perception.

### Control analysis: validating the effect of spatial arrangement with texture stimuli

Our results thus far suggest that what differentiates human perception from visual cortical and dCNN representations is sensitivity to spatial arrangement of features. However, an alternate explanation is that the superior performance of human observers at discriminating natural from synthesized images might be driven by low-level artifacts in the synthesis process for which dCNN observer models and high-level visual cortex are insensitive. To control for this potential confound, we utilized a class of stimuli, visual textures, which are inherently defined by their invariance to the spatial arrangement of features. If artifacts in the image synthesis process were responsible for the superior performance of human observers, then we would expect that human observers should still be much more likely than dCNN or cortical observer models to identify the natural texture image as the odd-one-out. However, if it is due to spatial arrangement as we hypothesized, then we would expect humans to be less able to accurately detect the natural texture image, resembling the performance of dCNN and visual cortical observer models.

We found that human observers’ oddity detection performance for texture images matched that of dCNN observer models. We selected 12 natural texture images (e.g. bricks, rocks, grass, moss, bark, etc.) with a relatively homogeneous spatial distribution of features and synthesized corresponding images which varied in their feature complexity and spatial arrangement. We compared the performance of human observers with five different dCNN observer models as we varied both the feature complexity and spatial arrangement of the synthesized images of textures. We found that human observers’ likelihood of identifying the natural image was not significantly different (*Linear mixed effects model: b=0.027, SE=0.014, p=0.066, 95% CI = [−0.002, 0.055])* from the dCNN models across all conditions (**Fig. S1A,B**). This finding, that human oddity detection performance resembles dCNN performance for textures but not for objects, provides evidence that what differentiates human perception from dCNNs is selectivity for spatial arrangement, not an artefact in the image synthesis process.

Using a representational geometry analysis, we found that the perceptual distance between natural texture images and their corresponding synthesized scrambled images only marginally exceeded (*t=3.38, p=0.006*) the perceptual distance between two different synthesized images with scrambled arrangements of features (**Fig. S1C**). In comparison to the perception of objects, the perceptual representation of textures was significantly less selective for natural feature arrangement (*t=-13.96, p<0.001).* Notably, the representational geometry of dCNNs (**Fig. S1D**) and human category-selective cortex (**Fig. S1E**) for textures also reflected a similar non-selectivity for the natural arrangement of features. In sum, these results suggest that the representations in human visual cortex and dCNN models can account for human perception of textures, which are invariant to feature arrangement, but cannot account for human perception of objects.

### Do neural and dCNN representations contain information about spatial arrangement?

To reconcile the discrepancy between neural representations and perception with regards to selectivity for natural feature arrangement, we sought to find a transformation of the cortical representation that might better approximate human perception. To quantify the ability of a particular representational space to distinguish natural objects from synths with scrambled matching features, we developed a natural image selectivity index that measures the degree to which the representational distance between the natural image and the synths exceeds the distance between two different synths.

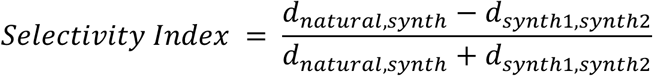

We found that an unweighted readout of the cortical representation yielded a natural image selectivity index that was not significantly different from zero (**Fig 7A**, purple bars) in nearly all category-selective visual areas (*mFus: t=1.21, p=0.27; pFus: 4.66, p=0.003; IOG: t=1.24, p=0.26; CoS: t=2.03, p=0.08; TOS: t=2.45, p=0.05; LO: t=0.97, p=0.37; VVC: t=2.09, p=0.08; PIT: t=-0.13, p=0.90; VMV: t=3.10, p=0.02),* with the marginal exception of pFus, TOS, and VMV. This contrasts sharply with the selectivity of perception (**Fig 7A**, dashed line), in line with our finding that the cortical observer model was unable to explain human behavioral performance (**Fig 4C**). We hypothesized that a linear transformation of cortical responses might yield a representational space that is more selective for the natural arrangement of object features. To test this hypothesis, using gradient descent, we sought to find a linear weighting of cortical responses which would yield a representation that was as selective for the natural arrangement of features as human perception.

**Figure 7.**
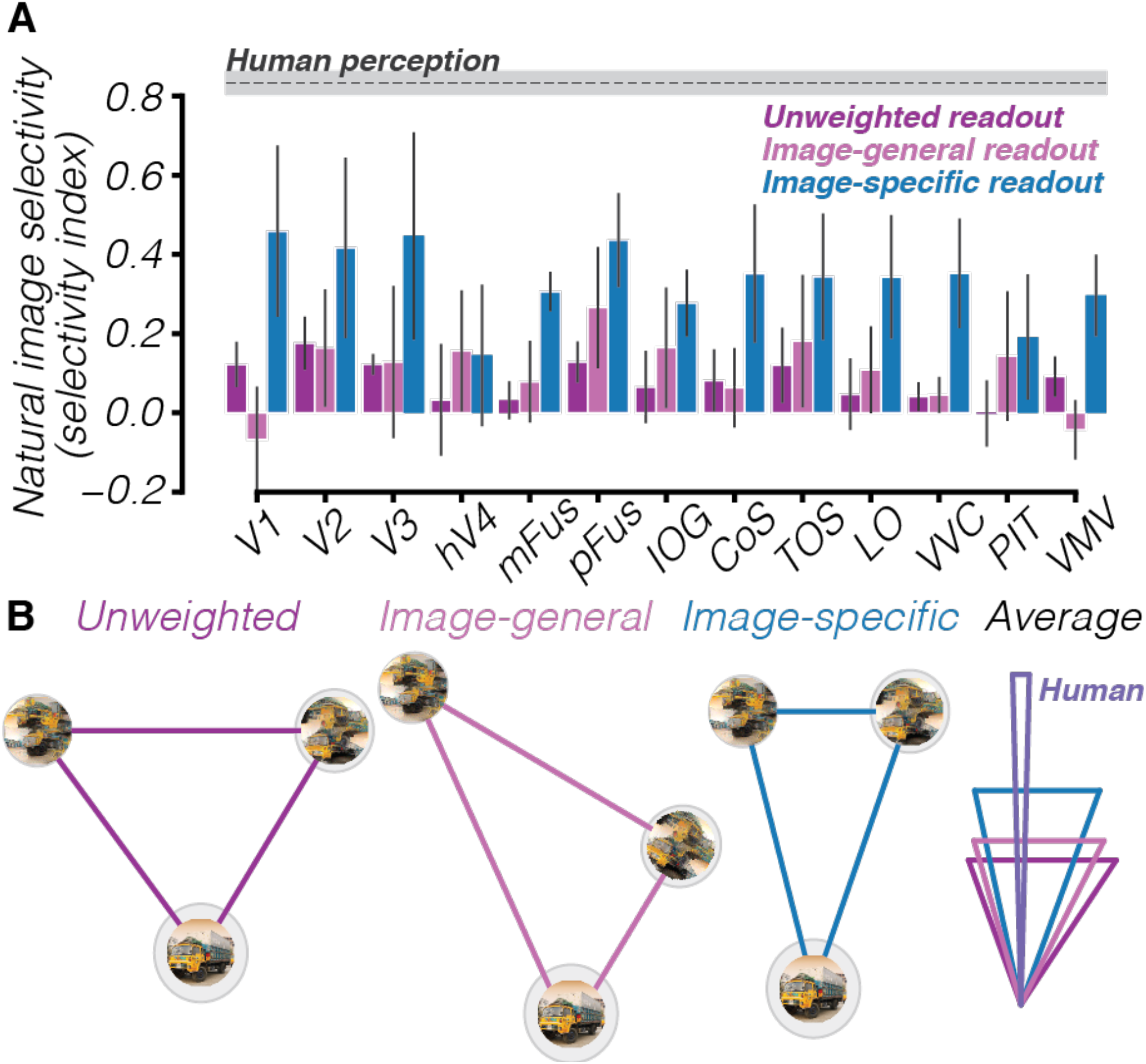
Evidence that information about natural feature arrangement can be read out from visual cortex. (A) Natural image selectivity index across all analyzed cortical regions, for unweighted readout (purple), image-general readout (magenta), and image-specific readout (blue). Gray dashed line shows natural image selectivity of human perception. (B) Representational geometry for one representative area (LO) for one example image class, comparing between unweighted, image-general, and image-specific readouts. Rightmost panel shows the representational geometry, averaged across all image classes.

We first tested a linear transform of visual cortical responses that generalized across image-classes but found that such a readout failed to increase selectivity. We used gradient descent to find a linear weighting of cortical responses that would maximize the natural feature arrangement selectivity index. To ensure that this transform would generalize, we cross-validated across image classes. That is, we fit the weights to maximize the natural image selectivity for all but one image class and then evaluated the selectivity of the weighted representation on the held-out image class. This procedure failed to yield a representation which was significantly more selective for natural feature arrangement than the unweighted readout in all regions of visual cortex (*V1: t=2.05, p=0.09; V2: t=0.23, p=0.82; V3: t=0.00, p=1.00; V4: t=-1.02, p=0.35; mFus: t=-0.75, p=0.48; pFus: t=-1.72, p=0.14; IOG: t=-1.09, p=0.32; CoS: t=0.56, p=0.59; TOS: t=-0.57, p=0.59; LO: t=-0.91, p=0.40; VVC: t=-0.69, p=0.52; PIT: t=-1.94, p=0.10; VMV: t=1.84, p=0.12)* (**Fig 7A,** magenta bars).

We next tested whether fitting a separate linear transform for each image class could increase the natural image selectivity of the representation. This would suggest that the representation at least contains enough information to extract the natural feature arrangement, even if it is not generalizable across image classes. To do so, we used gradient descent to find, for each image class, a weighting of cortical responses that would maximize the natural image selectivity index for that particular image class. Therefore, for each subject, within each brain area, we fit 1000 weights (100 voxels x 10 image classes). We cross-validated across presentations, such that weights were fit based on trial-averaged responses from 90% of the presentations and selectivity was assessed on the held-out 10% of image presentations.

We found that this image-specific weighting of cortical responses significantly increase the natural feature arrangement selectivity of the neural representation (*V1: t=-3.03, p=0.02; V2: t=-1.94, p=0.10; V3: t=-2.45, p=0.05; V4: t=-2.38, p=0.06; mFus: t=-12.94, p<0.001; pFus: t=-5.94, p=0.001; IOG: t=-3.63, p=0.01; CoS: t=-3.41, p=0.01; TOS: t=-2.74, p=0.03; LO: t=-4.85, p=0.004; VVC: t=-4.74, p=0.003; PIT: t=-2.81, p=0.03; VMV: t=-4.13, p=0.01)* (**Fig 7A,** blue bars) in all cortical areas, although this was still unable to reach the level of selectivity observed behaviorally (*V1: t=-2.99, p=0.024; V2: t=-3.00, p=0.024; V3: t=-2.53, p=0.045; V4: t=-7.34, p<0.001; mFus: t=-18.81, p<0.001;pFus: t=-6.79, p<0.001; IOG: t=-11.47, p<0.001; CoS: t=-4.65, p=0.003; TOS: t=-5.80, p=0.001; LO: t=-5.61, p=0.001; VVC: t=-6.40, p=0.001; PIT: t=-7.18, p<0.001; VMV: t=-8.55, p<0.001).* This form of readout, however, requires prior experience with every individual object to learn an image-specific linear transform, so it is unlikely to be a plausible mechanism by which visual cortical responses could directly support perception. Rather, it demonstrates that the information about natural feature arrangement can be found in these visual areas but would likely require further untangling to yield a representation which is more predictive of behavior.

Using a support vector machine (SVM) classifier trained to classify images as natural or synthesized, we found corroborating evidence that cortical responses contain sufficient information to distinguish natural from synthesized images, though not in a generalizable format. We trained a SVM classifier with a linear kernel on cortical responses from 13 different visual areas. When evaluated on samples from image classes within its training set, the classifier was highly accurate in classifying the sample as natural or synthesized (**Fig. S5A**, blue points). However, when evaluated on samples from image classes outside its training set, the classifier was unable to classify the images as natural or synthesized significantly above chance level, computed using a permutation test (**Fig. S5A**, magenta points). Thus, a linear classification boundary can be found that distinguishes natural from scrambled images, but the classification boundary varies for different image classes.

We found that a multilayer perceptron, which is capable of learning arbitrary nonlinear functions, can enhance the natural image selectivity of the dCNN feature representation. To test this, we analyzed the representational geometry of the last fully-connected layer in each deep convolutional neural network. We found that in 3 out of 4 dCNN model architectures, the last fully-connected layer was marginally more selective for natural feature arrangement compared to the last convolutional layer (*t=-2.98, p=0.058)* (**Fig. S6A**), although this fully-connected layer was still significantly less selective for natural feature arrangement compared to human observers (*t=-14.90, p<0.001).* This finding demonstrates the plausibility of a nonlinear readout to transform the feature representation in dCNNs or visual cortex to yield a representation which can better predict perceptual selectivity for natural feature arrangements of objects.

To address the possibility that information about natural feature arrangement is present in the dCNN representation but dominated by information about unlocalized features, we trained a support vector machine (SVM) classifier with a linear kernel to predict whether an image was natural or synthesized. We found that when the SVM classifier was evaluated on image classes that were not within its training set, it was unable to predict whether these images were natural or synthesized significantly above chance (**Fig. S2B**, magenta points), even if the SVM was trained exclusively on other image classes within the same category (**Fig. S2C**). However, when evaluated on image classes within its training set (**Fig. S2B**, blue points), the SVM classifier was able to predict whether an image was natural or synthesized. These results suggest that the representation of natural and synthesized images is sufficiently different that an image-specific classification boundary can be found but not an image-general boundary.

## Discussion

Through the measurement and analysis of human behavior, cortical responses, and dCNN features, we sought to characterize the visual representations which enable the perception of natural objects. We found that human visual perception is sensitive to the complexity of features and selective for the natural arrangement of object features. In contrast, we found that both dCNN features and ventral visual cortical responses were relatively poor at discriminating natural images from synthesized scrambles containing complex features. A model of macaque IT neurons was also similarly insensitive to the arrangement of features in object images. This insensitivity was not due to a lack of featural representations, but rather due to an insensitivity to spatial arrangement, as demonstrated by the evidence that representations in dCNNs, category-selective regions of human visual cortex, and macaque IT matched human performance at a category-level discrimination task and at a texture discrimination task, which did not require selectivity for feature arrangement. Thus, we concluded that both human visual cortex and dCNN models do not represent natural object images more distinctly than scrambled images of complex features that make up objects and are therefore unable to account for the human perceptual ability to discriminate natural from synthesized images of objects. This suggests that the ventral visual cortex is insufficient to fully support object perception (62) though it is a useful intermediate representation. To confirm this, we demonstrated that the information necessary to match perceptual selectivity for natural images is decodable from category-selective regions of human visual cortex, although it requires a specialized image-specific readout. Taken in sum, our results suggest that the representations found in human visual cortex, macaque IT cortex, and Imagenet-trained dCNNs encode the complex visual features that make up objects, though they do not distinctly encode the natural arrangement of features which defines an object.

Although the image synthesis technique that we employed allowed us to separately control the complexity and spatial arrangement of visual features in synthesized images, it is almost certainly the case that feature complexity and spatial arrangement are not fully independent dimensions. That is, complex visual features are composed of particular arrangements of simpler features. This is best exemplified by the recent finding that a dCNN with random filters at multiple spatial scales can be used to synthesize textures which are fairly perceptually similar to the original and comparable to the quality of textures synthesized by the Gatys model (75, 7). Nonetheless, it is useful to artificially vary the spatial arrangement of features at different levels of complexity to assess the contributions of each of these to perception. This utility is best demonstrated by the contrast between object and texture perception: whereas human observers’ perception of textures was sensitive only to the complexity of visual features and not to the spatial arrangement of those complex features, object perception was sensitive to both the complexity and arrangement of visual features. Further, this disentangling of feature complexity and spatial arrangement is justified by the finding that at a particular level of feature complexity (pool4), there was a mismatch between human perception and cortical representations in terms of selectivity for natural images relative to synthesized scrambled images.

To demonstrate the lack of cortical selectivity for natural feature arrangements, we used two convergent pieces of cortical evidence: BOLD imaging of human visual cortex and a model of macaque IT neurons. Although the BOLD imaging evidence was limited in its spatial and temporal resolution, it allowed us to measure neural responses in many visual areas across the occipital and temporal lobes. It is, however, possible that neural selectivity for natural feature arrangement is only observable at higher spatial (e.g. individual neurons) or temporal (e.g. individual spikes) resolutions. To attempt to address this concern in the absence of electrophysiological recordings, we utilized a well-validated model of macaque IT neurons. This method, too, had its limitations, as the model does not explain all the variance in IT neural responses, and it is possible that the unexplained variance might account for the perceptual selectivity for natural feature arrangement. However, even state-of-the-art techniques can only measure from a few thousand neurons in a localized region of the brain, which would leave open the possibility that selectivity for natural feature arrangements might be found when considering a larger, more representative sample of neurons. Therefore, given that both of our approaches, one with wide spatial coverage and another which models individual neuronal responses, yielded convergent results, we conclude there is a mismatch between human perception and representations in category selective cortex.

It is possible that this mismatch between human perception and cortical representations is due to subjects in the neuroimaging experiment passively viewing the images while performing a fixation task, in contrast to subjects in the behavioral experiment who were actively attending to the images. This possibility might imply that feedback, in the form of attention or other top-down signals, transforms visual cortical representations to facilitate the discrimination of natural from synthesized images. Therefore, our results should only be taken to apply to the feedforward cortical representations generated while passively viewing images. However, given that these passively generated feedforward representations were sufficient to match behavioral performance in the category-level oddity task, it is nonetheless noteworthy that feedforward responses do not preferentially represent natural images relative to synthesized scrambled images containing similar features.

Our results contribute to a large body of research about the texture-like encoding of peripheral vision (10, 18, 19). Despite the stimuli in our experiments being presented in the visual periphery, human subjects were highly sensitive to the spatial arrangement of features for object-like stimuli, although not for texture-like stimuli, in line with recent findings (14). We extend these findings to demonstrate that visual cortical representations of peripherally presented objects are not selective for the natural spatial arrangement of visual features. It is possible that had we measured cortical responses to foveally presented stimuli, we might have found a greater degree of selectivity for natural feature arrangement. However, the stimuli from the macaque IT dataset were presented at the center of gaze, yet we found that the model IT representation was insensitive to natural feature arrangement. Further, we presented the stimuli peripherally in the behavioral experiments while enforcing fixation (**Fig. S3**) and nonetheless observed selectivity for natural feature arrangement, so it is unlikely that our results could be explained by the texture-like encoding of peripheral vision alone.

These results suggest a possible cortical mechanism to explain a series of discrepant findings in the scene perception literature regarding why human observers cannot distinguish feature-matched synths (10) but can easily distinguish natural images of objects from feature-matched synths (14). An influential study (10) demonstrated that matching first and second-order statistics within spatial pooling regions, whose size matched V2 receptive fields, results in metameric images. However, subsequent work (14, 15) has shown that this metamerism only holds when comparing two synthesized samples but breaks down when one of the samples is the original image. It is particularly the case that human subjects can easily discriminate natural images from synths with scrambled features for objects or scenes more than for textures (14). If the synths are generated to match the locally pooled features of the natural image, then why might humans be able to discriminate the natural image from a synth but unable to discriminate two different synths? Our behavioral results corroborate the findings of (14), suggesting that image content – i.e. whether an image contains an object or a texture – matters beyond just the size of spatial pooling regions in terms of whether a natural image is perceptually distinct from synthesized counterparts. Our BOLD imaging data suggest a possible cortical mechanism: a specialized object-specific readout from visual cortex can support the enhanced discriminability of natural objects from synths.

In the domain of face perception, prior research has sought to distinguish selectivity for complex features from selectivity for the arrangement of those features. There is evidence suggesting that face-selective visual areas in the ventral temporal cortex are sensitive to the spatial arrangement of facial features, as demonstrated by the finding that the mean response of all voxels in the fusiform face area is greater to intact faces than faces containing scrambled facial features (60), as well as the finding that a linear classifier can decode intact vs. scrambled faces (61), which might seem to contradict our findings. However, these prior findings relied on handpicked features or grid-based scrambling approaches. Due to the limited number of stimuli in our sample, we did not specifically examine selectivity for spatial arrangement of facial features, so we cannot make any claims about whether our approach using deep image synthesis would confirm or contradict prior studies using handpicked features. Regardless, given the degree to which face-selective cortical populations appear to be highly functionally specialized and anatomically segregated compared to other object-selective populations in VTC and LO, it is certainly possible that the neural representation of faces might be explicitly selective for the natural arrangement of facial features, while most other objects are represented by a collection of disjointed complex visual features.

Our results contribute to a long-standing debate about whether the perception of objects is holistic or featural. Behavioral effects, such as the Thatcher effect (76), in which subjects are relatively insensitive to local feature rotations in an upside-down face, or the finding that humans are better at identifying facial features when presented in context than in isolation (77, 78), have given rise to the view that objects must be represented holistically, not as an independent set of features. In the present study, we assess holistic perception as the degree to which the whole object is perceptually distinct from the scrambled features of the object. Using the oddity detection task, we found that the natural image is far more perceptually distinct from a synthesized scrambled image than two different scrambled images are from each other, suggesting that humans perceive objects holistically. However, cortically, we found that natural object are not represented more distinctly from synths than two different synths are from one another, suggesting a non-holistic representation. These results suggest that a holistic perception of objects may arise from a featural cortical representation.

Our findings build upon recent research demonstrating that Imagenet-trained deep convolutional neural network models are texture-biased, i.e. more likely to make use of texture than shape information when classifying images, in contrast to humans who are shape-biased (53, 54, 56, 79). However, while these prior studies were taken as evidence that dCNNs are a poor model of visual representations in the human brain, our results demonstrate that representations in the visual cortex are as texture-like as those of dCNNs and therefore both dCNNs and category-selective regions of visual cortex similarly deviate from human behavior. When texture and shape cues are artificially made to conflict, either by using silhouetting (54) or by using neural style transfer (53), Imagenet-trained dCNN models are more likely to classify the conflicting images according to the texture than the shape label, unlike humans. It is possible that the texture bias of dCNNs is driven by the readout rather than the feature representations themselves. Our results demonstrate that dCNNs’ texture-bias is driven by the lack of selectivity for natural spatial arrangement in the feature representation itself, not by a texture-biased readout. We note that the term shape, while related to the concept of feature arrangement, is often used to refer to external contour. While our image synthesis technique does not specifically target external contours, it does disrupt them along with any other natural arrangement of features. Finally, our findings examining cortical representations of objects suggest a reinterpretation of the texture-bias results (53, 54). In contrast to the suggestion that the texture bias of dCNN models makes them flawed as models of human vision, we found that the human visual cortex similarly contains texture-like representations of objects. We can thus speculate that this texture-like representation, which is non-selective for natural feature arrangements of objects, might be useful for the perception of objects. This might seemingly contradict the finding that texture-biased dCNNs are more susceptible to image distortions than their shape-biased counterparts (53). However, our results suggest the possibility that texture-like representations need not necessarily be less robust, if coupled with a sufficiently sophisticated readout.

What potential advantage is there to a representation that encodes complex visual features but does not prioritize the natural arrangement of these features? Classic theories of object vision have posited that objects can be identified by the arrangement of more simple building blocks or primitives (43). Our findings suggest that textures are one of these basic building blocks that are encoded by category selective cortex. Category selectivity of visual cortical areas then comes from visual features that are informative for particular category judgements. For example, parahippocampal place area, though known for selectivity for scenes and places (80, 81), is also highly active for more simple rectilinear features that are building blocks for that category (82). Our findings further show that despite not being selective for the natural arrangement of visual features, the natural arrangement of complex features has not been lost in the cortical representation. That is, a classification analysis could decode (on an item-by-item basis) the natural spatial arrangement. If sensory representations were instead specific for a particular arrangement of complex visual features, this might preclude the possibility of learning new arrangements of features for novel object categories. The implication is thus that it is beneficial to have a feature representation in high-level visual cortex that is non-selective for natural spatial arrangement because it might allow for more rapid, robust transfer learning: that is, the repurposing of learned features for novel objects or tasks. This is certainly true of Imagenet-trained dCNN models, whose learned feature space can be easily repurposed for new object categories or even for other tasks (83, 84), simply by learning a new linear readout. Taken together, our results suggest that cortical visual responses in category-selective regions represent not objects per se, but a basis set of complex visual features that can be infinitely transformed into spatial arrangement sensitive representations of the myriad objects and scenes that we encounter in our visual environments.

## Materials and Methods

### Behavioral Methods

#### Observers

87 observers, naive to the goals of this study, performed 100 trials of the oddity detection behavioral experiments and 110 observers performed 100 trials of the dissimilarity judgment experiment, on Amazon Mechanical Turk. Subjects were eligible to participate if their HIT approval rate (percentage of completed HITs that were approved by prior requesters) exceeded 75%. Subjects’ data were excluded only if their performance on trivial catch trials failed to exceed 40% accuracy (chance level performance was 33%). To validate the online findings, in-lab behavioral data were collected from 2 observers, where each observer participated in at least 5000 trials. For the neuroimaging experiment, seven observers (4 female, 3 male, mean age 28 y, age range 23 - 37 y), 6 naive to the goals of the study, participated as subjects in two 1 - hour scans at the Stanford Center for Neurobiological Imaging. All protocols were approved beforehand by the Institutional Review Board for research on human subjects at Stanford University, and all observers gave informed consent prior to the start of the experiment by signing a form.

#### Stimulus Generation

The stimuli used in all experiments were either natural images of real objects/textures or synthetically generated through an iterative optimization procedure (“synths”) designed to match features from a target image. To synthesize images, we passed a target natural image into an Imagenet-trained VGG-19 dCNN model (63) and extracted the activations from 3 intermediate layers (pool1, pool2, pool4) (**Fig 1A**). We calculated a spatially constrained Gramian matrix, i.e. the inner product between every pair of activation maps, of each layer’s activations, which allowed us to spatially pool features over image subregions of pre-defined sizes. Finally, we iteratively updated the pixels of a random white noise image to minimize the mean squared error between the spatially weighted Gramians of the output image and the target image. Thus, by varying the size of the spatial pooling regions, we could vary the degree to which feature arrangement was spatially constrained, and by varying which layers’ features were included in the loss function of the optimization, we could control the complexity of visual features in the synthesized image. See extended methods for more detail.

#### Experimental Design - Oddity Detection Task

On each trial (**Fig 2A**), observers were concurrently presented with 3 images (1 natural, 2 synths) for up to 2 seconds and asked to select the one which appeared most different from the others. Both synths were matched to the features of the natural image, at a particular dCNN layer and spatial constraint level. We performed the experiment both on Amazon Mechanical Turk (N=87 subjects, 6165 trials) and in the lab (N=2 subjects, 5000 trials each). See extended methods for more detail.

#### Estimating perceptual distances

Using behavioral responses from the pairwise dissimilarity judgment task, we estimated the perceptual distances between pairs of images. We used the maximum-likelihood distance scaling model (73) to estimate the perceptual distances between pairs of images as free parameters in an optimization procedure designed to maximize the likelihood of the observed behavioral responses. See extended methods for more detail.

### dCNN Methods

We modeled task performance using features extracted from a deep convolutional neural network (dCNN). We tested the representational space of five different Imagenet-trained dCNNs: VGG-19 (63), CORnet-Z (64), VGG-16 (63), ResNet-18 (65), and AlexNet (46). On each trial, our model extracted a feature vector from the last convolutional layer of the dCNN for each image presented (**Fig 3A**). Next, we computed the Pearson distance between the features of each pair of images, and for each image, calculated its dissimilarity as the mean Pearson distance between that image and each of the other two images. Finally, the model converted these dissimilarities into choice probabilities using a Softmax transform. See extended methods for more detail.

To analyze the representational space learned by Imagenet-trained dCNN models, we employed a representational similarity analysis to compute the Pearson correlation distance between the activations from the last convolutional layer in response to each pair of images (85). We then used these representational distances to determine the selectivity of this feature space for natural feature arrangement, computed as the difference between the *d_natural,synth_* and the *d_synth,synth_*, normalized across categories.

#### Modeling IT neurons

To estimate object selectivity of neurons in inferior temporal (IT) cortex, we modeled the response of each IT neural site (71) as a linear combination of dCNN units (48, 51, 86, 87). Using this population of model IT neurons, we modeled oddity detection task performance and calculated the representational selectivity for natural feature arrangement. See extended methods for more detail.

### Neuroimaging Methods

We used blood-oxygen level dependent (BOLD) imaging (92) to measure cortical responses to visually presented images, both natural and synthesized (1×1 pool4), from 10 different categories. Images subtended 12 degrees. We defined cortical areas using population receptive field mapping to identify retinotopic areas in early visual cortex (68, 88–90), in addition to an atlas-based approach to identify anatomically defined areas (70) and a functional localizer to identify category-selective regions (69). Using a generalized linear model (66, 67), we extracted trial-averaged responses to individual images. We used these responses as input to observer models to perform the task as well as to compute the selectivity of neural representations for natural feature arrangement. See extended methods (Neuroimaging Methods section) for more details.

## Acknowledgments

We acknowledge the support of Research to Prevent Blindness and Lions Clubs International Foundation and the Hellman Fellows Fund to J.L.G. This work was supported by the Stanford Center for Cognitive and Neurobiological Imaging.

