## Supplemental Information for "Texture-like representation of objects in human visual cortex"

**This PDF file includes:**

Supplementary text (extended methods)
Figure S1: Control analyses using texture stimuli
Figure S2: SVM classification accuracy in visual cortex and dCNNs
Figure S3: Behavior comparison in-lab vs online
Figure S4: Behavior comparison with vs without feedback
Figure S5: Pairwise dissimilarity task behavior
Figure S6: Natural image selectivity of dCNNs
Figure S7: All stimuli used in neuroimaging experiment
Figure S8: Synthesized images at different iterations

References for supplementary text

### Extended methods

#### ***Stimulus generation***

All stimuli used in these experiments were either natural images, drawn from image searches of Creative Commons licensed texture and object images, or were synthetically generated through an iterative optimization procedure (“synths”). We selected 34 natural images (22 objects and 12 textures), cropped them into squares, and downsampled the images to 256 x 256 pixels. Images were selected as “object stimuli” only if they contained exactly one object, either animate or inanimate, that was clearly visible in the image. Images were selected as texture stimuli if they contained repeated patterns and/or numerous objects that were similar in appearance. Images were categorized as textures or objects by the authors prior to data collection, and these category judgments were confirmed using a Mechanical Turk experiment where an independent sample of 65 subjects were asked to categorize each image as either a texture or object (mean correlation between subjects’ categorizations and our categorization was 0.924).

The image generation procedure, adapted from (1) involved three major steps: feature extraction, spatial pooling, and image synthesis via pixel-wise optimization. In the feature extraction stage, each natural image was passed into an Imagenet-trained VGG-19 deep convolutional neural network (2), and the activations of 3 intermediate layers (pool1, pool2, and pool4) were extracted (Fig 1a). The spatial pooling stage of the standard Gatys algorithm was done by computing the Gramian matrix, i.e. the inner product between pair of activation maps, in each layer. The Gramian matrix preserves individual features as well as the co-incidence of features, while discarding information about the spatial position of those features. Finally, using gradient descent with the L-BFGS algorithmic solver (3), we updated the pixels of a random white noise image to minimize the mean squared error to the Gramian computed for the natural image (1, 4, 5).

This image synthesis algorithm allowed us to control the complexity of the features in the natural image which are matched in the synthesized output. By varying which layers were included in the loss function, we controlled the complexity of the features in the generated image. This is based on prior research suggesting that early layers of dCNNs encode simple features, such as orientation and spatial frequency, whereas later layers encode more complex features, such as texture, shape, or category identity (6–10). This is an improvement over other texture synthesis algorithms, such as the Portilla-Simoncelli algorithm (4), which includes only one level of higher order statistics computed from the pairwise correlations of a V1-like filterbank (11). We selected 3 different layers from the VGG19 model to include in the synthesis procedure: pool1, an early layer (64 filters); pool2, an intermediate layer (128 filters); and pool4, a late layer (256 filters). Layers were added incrementally, so images generated in the pool1 condition include only pool1 features, whereas images generated in the pool4 condition include features from layers pool1, pool2, and pool4.

This image synthesis algorithm also allowed control over the spatial scale within which the spatial arrangement of features is constrained. Whereas the original Gatys algorithm (1) pools features across the entire image, we modified the algorithm to compute spatially weighted Gramians, which only pooled features within pre-defined spatial pooling regions (4). We tiled the image with equal-sized square spatial pooling regions with smooth transition boundaries defined by a squared cosine function with 20 pixel ramping boundaries. For each unit in the model, we calculated the overlap between its receptive field and each spatial pooling region. We used this to compute a spatially weighted Gramian matrix for each pooling region, wherein units are included in proportion to how much their receptive field overlaps the spatial pooling region. By varying the size and number of spatial pooling regions, we imposed stronger or weaker constraints on the spatial arrangement of features. Low spatial constraint synths are ones in which the spatial arrangement of the features can be scrambled across the entire image (which we call 1x1 as there is a single spatial pooling region) and high spatial constraint synths are those in which the arrangement of the features are constrained within small subregions of the image, (for example, a

4x4 set of spatial pooling regions constrains features in subregions that are 1/16th the area of the full image).

The number of parameters that constrained each image was a function of the size of the Gramian and the number of spatial pooling regions. The size of the Gramian for a particular layer is equal to the square of the number of filters in each layer, so pool1 images were constrained by 4096 ($64^2$) parameters, pool2 images were constrained by 20480 parameters ($128^2 + 64^2$), and pool4 images were constrained by 282624 parameters ($512^2 + 128^2 + 64^2$). To constrain spatial arrangement of features, we computed a separate Gramian for each spatial pooling region, so the number of spatial pooling regions was a multiplier on the number of parameters. For example, the 4x4 images contained 16x the number of parameters as the 1x1 images.

Finally, by initiating the optimization process with different random seed images, we generated multiple different synthesis samples which differed significantly in the pixel representation space but contained nearly identical features within each spatial pooling region. For each natural image, we synthesized 3 samples at each of 3 layers (pool1, pool2, and pool4) and 4 spatial constraints (1x1, 2x2, 3x3, 4x4), for a total of 36 synthesized samples per natural image. We used the Adam optimizer (12), implemented in Tensorflow (13), and terminated the image synthesis optimization after 10,000 iterations.

### ***Behavioral Methods***

#### *Experimental Design – Oddity detection task*

In the oddity detection experiment, observers performed a 3 alternative forced choice judgment of the odd-one-out (14). On each trial (**Fig 2A**), observers were asked to fixate centrally on a cross for the duration of the trial, although we could not enforce fixation with eye-tracking and did not employ a central task at fixation. After 200ms of fixation, 3 images were presented -- 1 natural and 2 synths -- concurrently for 2 seconds. Observers were instructed to respond within 2 seconds, using a keypress, to indicate which image was most different from the others, and on 89.04% of trials, subjects did respond within the time limit (mean RT: 1.08s, SD=0.41s). The two synths were always generated to match the features of the natural image and were both generated from the same layer and spatial constraint, with a different random seed. Following the subject's response, they were shown feedback in the form of the fixation cross changing color for 200ms to either green, indicating a correct response, or red, indicating an incorrect or no response. On each trial, we randomly selected a natural image and 2 different synthesized samples, both with the same feature complexity and spatial constraints, to display. Images subtended approximately 8 degrees, though there was some variability due to the screen and window size of individual participants. Each image was centered 6 degrees away from the fixation cross. We performed this experiment both on Amazon Mechanical Turk, where we recruited 87 subjects who performed a total of 6165 trials, as well as in the lab, where we recruited 2 subjects to perform a total of approximately 5000 trials each and were able to enforce fixation using an Eyelink eyetracking system that aborted any trials where subjects' eye-gaze deviated more than 1 degree from the fixation cross. A comparison of in-lab and online data is presented in Supp. Fig. S3. We presented 34 different image classes in this behavioral experiment, including 22 object image classes (Fig. 2) and 12 texture image classes (Supp. Fig. S1), where an image class is defined as the set of images including a natural image and all corresponding feature-matched synthesized samples.

#### *Experimental Design - Pairwise dissimilarity judgment task*

To determine the perceptual similarity of synths with naturals, we conducted a dissimilarity judgment experiment with an independent set of 110 observers. On each trial, observers were shown 4 images, grouped into two pairs and were asked to indicate with a keypress which of the two pairs was more dissimilar. Images subtended 8 degrees. Each pair was centered 8 degrees to the left and right of fixation, with 4 degrees of vertical separation between each image. Subjects fixated for 200ms and then stimuli were presented for 2 seconds and subjects were allowed to respond any time before the images disappeared. No feedback was given. As with the oddity detection task, all images presented on a given trial were generated to match the same natural image, and all synths were of the same feature complexity and spatial constraint. However, unlike the oddity detection task, on a randomly interleaved half of all trials, all 4 images were synths, and on the other half of the trials, 1 image was the natural image and the other 3 were synthesized images with scrambled arrangements of features. This enabled us to determine the perceptual similarity between the synths and the naturals as well as the perceptual similarity between different synths. We average together all of the distances between pairs of synths to yield a single synth-synth distance. Across all trials, subjects saw 170 unique images: 34 image classes x (1 natural image + 4 synthesized images). We collected a total of 8687 trials across 110 observers.

#### ***Estimating perceptual distances***

On any given trial, the observer saw 4 images grouped into 2 pairs, $(i_1, i_2)$ and $(i_3, i_4)$, and was asked to report which pair was more dissimilar. We can thus represent the probability that the observer will select the first pair $(i_1, i_2)$ as:

$$\mathrm{P}\left(D_{1,2} - D_{3,4} + \epsilon > 0\right)$$

where $D_{1,2}$ represents the perceptual distance between the first pair of images, $D_{3,4}$ represents the distance between the second pair of images, and $\epsilon$ is a Gaussian-distributed random variable with mean 0 and standard deviation $\sigma$ representing the combination of sensory and response noise. Then, the probability that the observer will select the first pair is given by $P\left(\epsilon < D_{1,2} - D_{3,4}\right)$, which can be computed as the cumulative distribution function of $\epsilon$, $\Phi(x)$ evaluated at $D_{1,2} - D_{3,4}$. The probability of selecting the second pair is then given by $1 - \Phi\left(D_{1,2} - D_{3,4}\right)$. Over $N$ trials, if we observe responses $r_1, \ldots, r_N$, we can compute the likelihood of observing these responses given the pairwise distances, as

$$P\left(r_1, \ldots, r_N \middle| D_{1,2}, D_{1,3}, \ldots, D_{N-1,N}, \sigma\right) = \prod_{i=1}^{N} \Phi\left(D_{i_1,i_2} - D_{i_3,i_4}\right)^{r_i} \times \left(1 - \Phi\left(D_{i_1,i_2} - D_{i_3,i_4}\right)\right)^{1-r_i}$$

Then, we used the Nelder-Mead optimization algorithm, as implemented in the Python scipy library (15), to find the values of the distances and the $\sigma$ that maximize this likelihood function. For each of the 34 image classes (which were also presented in the oddity task), we estimated the pairwise distances between 5 images (1 natural, 4 synth), resulting in 10 pairwise distances ($\binom{5}{2}$) to estimate for each image class, yielding a total of 341 parameters (including $\sigma$) that were estimated on 8687 trials of oddity detection behavior.

#### ***dCNN observer model***

On each trial, our model extracted a feature vector from the last convolutional layer of the dCNN for each image presented (**Fig 2B**). Next, we computed the Pearson distance between the features of each pair of images, and for each image, calculated its dissimilarity as the mean Pearson distance from the other two images. Finally, the model converted these dissimilarities into choice probabilities using a Softmax transform. Thus, the probability of choosing the i$^{th}$ item is given by:

$$P(c_i) = \frac{e^{\beta c_i}}{\sum_{j=1}^{3} e^{\beta c_j}},$$

where $\beta$ is the only estimated parameter, shared across all trials, image classes, and subjects, that is fit to maximize the likelihood of the observed choices and $c_i$ is the mean distance of the i$^{th}$ image from the other two. The $\beta$ parameter controls the extent to which the model maximizes the choice probability of the most dissimilar image, where a $\beta$ of 0 yields equal choice probabilities for all images and a beta of infinity would result in a choice probability of 1 for the most dissimilar image. We can then visualize these trial-by-trial choice probabilities by computing the average across all the trials of a single condition, to compare the behavior of the model to that of the human subjects.

***Modeling IT neurons***

To assess the selectivity of neurons in inferior temporal (IT) cortex for natural feature arrangement, we fit a model to a published dataset (16) of multielectrode array recordings measured while macaques passively viewed images of various objects serially presented at the center of gaze. We estimated the response of each neuron as a linear function of activations from each layer of an Imagenet-trained deep convolutional neural network (17, 18), a well-validated approach which yields state-of-the-art predictions of IT neural responses (19, 20). By finding the optimal weighting of dCNN features for best predicting each IT neuron's response, we could then compute a prediction of how each IT neuron would respond to novel images. Then, using this population of 168 model IT neurons, we computed the Pearson distance between the model population's response to each natural image and a corresponding synthesized image as well as the Pearson distance between the model population's response to two different synthesized images of the same class (**Fig 5A**). Using these two distance measures, we were able to compute a normalized index of selectivity for natural feature arrangement by the formula:

$$\frac{d_{natural,synth} - d_{synth1,synth2}}{d_{natural,synth} + d_{synth1,synth2}}$$

Given that our IT model explains, on average, 51.8% of the cross-validated variance in IT neural responses to naturalistic images (18), we cannot treat this as a perfect approximation of IT neurons, although we can use this as a reasonable proxy for IT single unit responses, to corroborate our BOLD imaging evidence. (See Discussion for further consideration of the caveats of this modeling approach).

***Neuroimaging Methods***

***BOLD Imaging Data Collection***

To measure neural responses to natural and synthesized images, we conducted an experiment using blood-oxygen level dependent (BOLD) imaging (21). We recruited seven subjects and instructed them to fixate while visual stimuli were presented over the course of two sessions. To identify the retinotopic maps in visual cortex (22, 23), we presented subjects with four 4-minute runs of a high-contrast sweeping bar stimulus, while they performed a color discrimination task at the center of the screen to ensure fixation (24). To identify and map category-selective regions in the ventral temporal cortex, we presented subjects with four 5-minute runs in which stimuli drawn from 5 categories (characters, bodies, faces, places, objects) were presented in a block design, while subjects performed a 1-back working memory task (25). Finally, to compare the neural response to natural images to their synthesized scrambled counterparts, we presented subjects with at least eight 6-minute runs, in which images were presented for 4 seconds, with no interstimulus interval, in an event-related design (26, 27), while subjects performed a central fixation task. Images subtended 12 degrees and were presented on both the left and right sides of the screen, centered at an eccentricity of 7 degrees. We selected 10 different image classes, consisting of 7 objects and 3 textures, and for each class, presented 1 natural image and 2 synthesized images, generated at a spatial constraint of 1x1 and from the pool4 layer. We also matched the Fourier magnitude spectrum and the luminance histogram of the synthesized images to their corresponding natural image, to control for potential low-level confounds. Over the course of the entire experiment, each image was repeated approximately 20 to 24 times.

All scans were collected on a 3 Tesla General Electric MRI scanner, using a T2* weighted sequence with multiplex factor of 4 (13 slices at multiplex 4 = 52 slices total), voxel size of 2.5mm, repetition time (TR) of 1.0s and echo time (TE) of 30ms. Additionally, we acquired a whole-brain high-resolution T1-weighted 3D BRAVO sequence with 0.9mm isotropic voxels. This anatomical image was used for segmentation and surface reconstruction, which were performed using Freesurfer. To correct for susceptibility distortions, we acquired an additional T2* weighted sequence with reversed phase encoding direction and used the TOPUP function from FSL (28).

We performed volume-by-volume image registration to correct for motion artefacts using standard procedures for motion correction (29). In the second session, we acquired another T1-weighted 3D BRAVO scan with voxel size 1.2 x 1.2 x 0.9mm. Using an image-based registration algorithm (29), we aligned this anatomical scan to the high-resolution anatomical scan so that functional regions of interest defined from the first session could be used to analyze the second session's functional data.

***Defining cortical areas***

Using a 3-parameter population receptive field (pRF) model, we estimated the center (x,y) and width (sigma) of the receptive field of each voxel in the occipital lobe (22). Then, we manually drew visual area boundaries delineated by the reversal in the gradient of the polar angle of pRFs (30). We were able to identify V1, V2, V3, and hV4 in all 7 subjects.

To identify category selective visual areas, we used the fLoc functional localizer (25), in which images of faces, bodies, places, characters, objects, and phase-scrambles were presented in a block design. We then used a GLM to estimate the response amplitudes to each stimulus category and then performed a statistical contrast to identify category-selective voxels. We were able to identify 3 face-selective clusters of voxels, in the mid-fusiform sulcus (mFus), posterior fusiform gyrus (pFus) and inferior occipital gyrus (IOG) and 2 place-selective clusters of voxels, in the transverse occipital sulcus (TOS) and the collateral sulcus (CoS), in each subject. We also used an atlas-based approach to identify anatomically defined areas (31), using a surface-based alignment to align the atlas to each subject's individual brain. We analyzed responses in 4 visual areas from the Glasser Atlas: lateral occipital complex (LO), for which we combined 3 smaller subregions, LO1, LO2, and LO3; ventral visual cortex (VVC); posterior inferotemporal cortex (PIT); and ventromedial visual area (VMV) for which we combined VMV1, VMV2, and VMV3 (31). These areas were selected because they have been identified as regions that contain information about visual object category.

In all analyses, we thus examined a total of 13 visual areas: 4 retinotopically defined areas (V1, V2, V3, hV4), 5 category-selective areas defined by a functional localizer (mFus, pFus, IOG, TOS, CoS), and 4 anatomically defined areas from the Glasser atlas (LO, VVC, PIT, VMV).

***BOLD Data Analysis***

We extracted trial-averaged neural responses to individual images using the GLMdenoise Matlab package (27), which estimates noise regressors from task-irrelevant voxels and uses those in a generalized linear model (GLM) (32). To identify the most reliable voxels in each cortical area, we split the data into two sets, such that each set contained half of the trials in which a particular image was presented. Then, we re-fit the GLM separately to each of the two splits and for each voxel, computed the correlation between its responses across the two splits. This measure of split-half correlation was used to identify the most reliable voxels, and in all analyses, we selected the 100 most reliable voxels in each ROI.

To determine how selective each visually responsive region is for the particular spatial arrangement of features that is found in the natural image, we computed the Pearson distance between the cortical response to a natural image and the neural response to a synth of the same class ($d_{natural,synth}$). We also computed the Pearson distance between the cortical response to two different synths of the same class ($d_{synth,synth}$) (**Fig 4A**). Finally, we computed the average Pearson distance between the cortical response to a synth of one class and the synths of every other class and called this the "between-class" distance. We assessed the category selectivity of a given cortical population by the degree to which the between-class distance exceeded the within class distance.

***<u>Triangle plot visualization</u>***

To visualize the relative representational distances between pairs of images, we plotted images in a triangle, where the length of the edges represents the magnitude of the representational distance between that pair of images. The representational distances are computed as the Pearson distance between each pair of images, for the dCNNs, cortical responses, and the model IT responses, but are estimated using maximum likelihood estimation for the human perceptual distances. Given the distances between 3 images, it is always possible to create a triangle where the edges correspond to distances, as long as none of the edge lengths exceeds the sum of the other two edge lengths. Then we rotate and translate the triangle so that the oddity image is always placed at the origin and the non-oddity images are above the oddity.

### ***Readout analyses***

#### ***Selectivity index***

We quantified the selectivity for natural feature arrangement by the degree to which the natural-synth distance exceeded the synth-synth distance, normalized by the sum of the natural-synth distance and the synth-synth distance.

$$Selectivity\ Index = \frac{d_{natural,synth} - d_{synth1,synth2}}{d_{natural,synth} + d_{synth1,synth2}}$$

This measure reflects the extent to which a representation differentiates the natural image, i.e. the extent to which the natural image is more different from the synthesized images than the synthesized images are from each other.

#### ***Image-general readout***

We performed an image-general readout by fitting weights to each voxel with the objective of maximizing the selectivity index across all image classes. We tested for generalization by fitting the weights to maximize the selectivity index for all but one image class and then evaluating the selectivity index on the held-out image class. Therefore, the number of parameters was equal to 100 (number of voxels) per area.

#### ***Image-specific readout***

We performed an image class-specific readout by fitting a separate set of weights to each voxel for each image class, with the objective of maximizing the selectivity index for each image class separately. Therefore, this approach required 1000 parameters per visual area (10 voxels x 10 images). To prevent overfitting, we estimated betas for each trial separately, then randomly selected 90% of the trials, averaged together the betas, and fit the weights on that 90% of trials for each image class separately. Then we evaluated the selectivity index on the held-out 10% of trials. We selected 100 voxels for inclusion in this analysis by separately splitting up the 90% of trials into two halves and choosing the voxels which had the highest split-half reliability in this subset of the data. This approach therefore ensured that no part of the weight estimation could be influenced by the held-out trials.

### Supplementary Figures

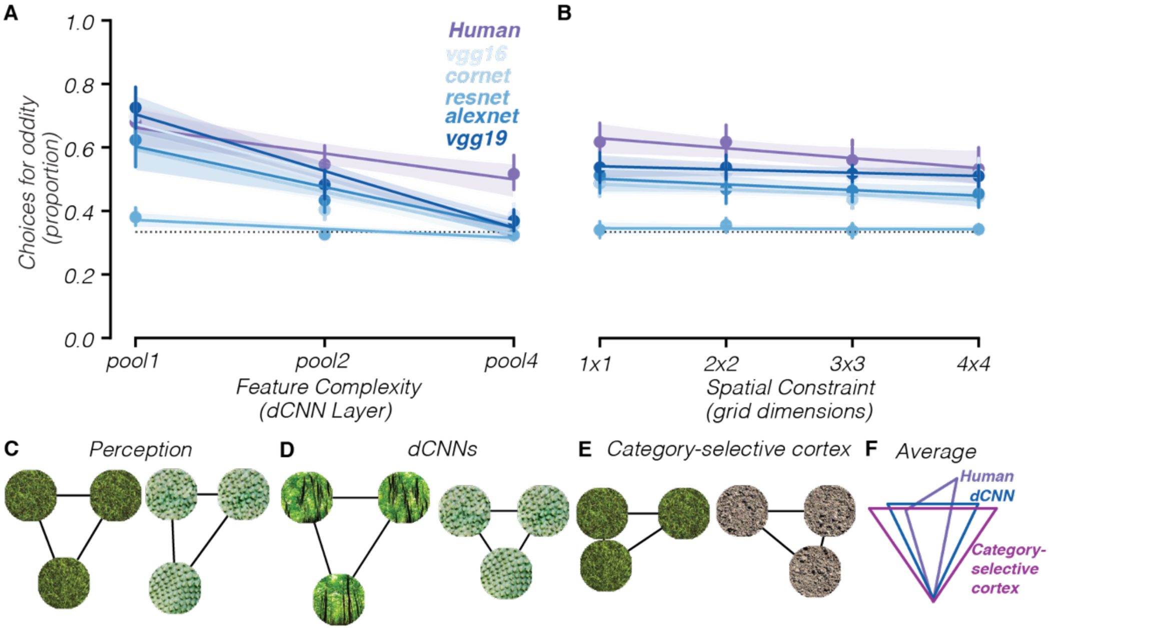


**Supp. Fig. S1.** Human observers are less sensitive to natural feature arrangement for texture-like images, similar to dCNN observer models and VTC voxels. (A) Performance of human observers (purple) compared to dCNN observer models (blues) at identifying the natural image as a function of feature complexity of synthesized images. (B) Performance of human observers (purple) compared to dCNN observer models (blues) at identifying the natural image as a function of constraints on spatial arrangements. (C-E) Triangular distance plots for perception (C), dCNN observer models (D), and category-selective cortical areas (E). (F) Average triangular distance plot across image categories, comparing category-selective cortex (magenta), dCNNs (blue), human observers (purple).

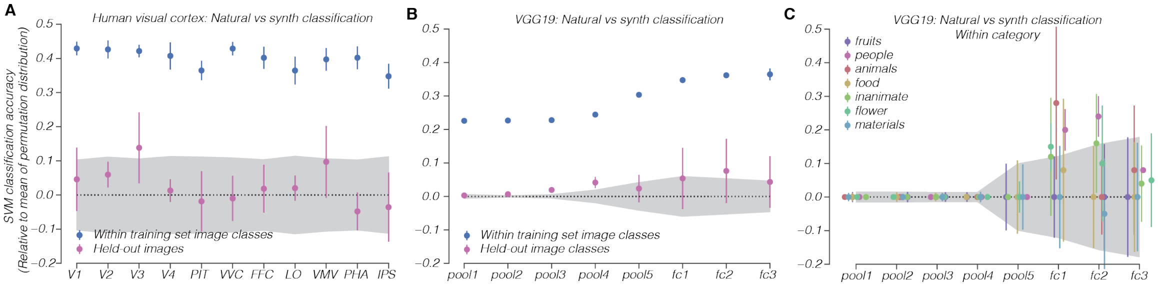


**Supp. Fig. S2**. Classification accuracy using support vector machines to classify natural vs synth. (A) Human visual cortex natural-vs-synth classification accuracy, relative to mean of permutation distribution. Gray shaded region represents 95% confidence interval of permutation distribution. Blue points are classification accuracy for image classes within the training set, and pink points are classification accuracy for image classes that were not in the training set. (B) Same as A but using features from various VGG19 layers instead of cortical responses. (C) Within-category decoding accuracy. We grouped 37 image classes into 7 categories and trained a SVM classifier to predict whether an image was natural or synthesized on all image classes of the same category except one and evaluated its performance on the held out image class of the same category. Across 7 categories (fruits, people, animals, food, flowers, inanimate objects, and materials), we found that classification accuracy failed to exceed chance in layers pool1, pool2, pool3, pool4, and pool5, although classification accuracy did exceed chance level for two categories (animals, people) in fc1 and one category (people) in fc2.

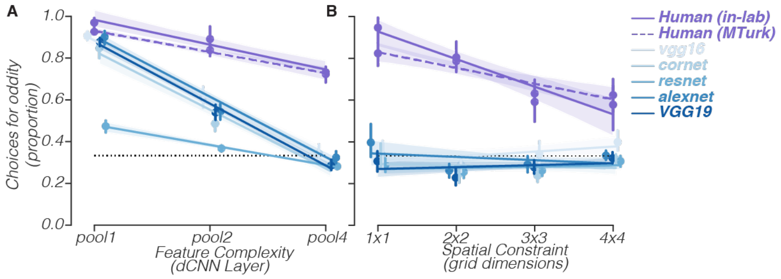


**Supp. Fig. S3.** Replication of behavioral results using dataset collected in-lab where fixation could be enforced with eye-tracking. (A) Comparison of human and dCNN behavior as a function of feature complexity. Solid purple line represents in-lab data and dashed purple line represents online data. (B) Comparison of human and dCNN behavior as a function of spatial constraint, fixing the feature complexity at the highest level (pool4).

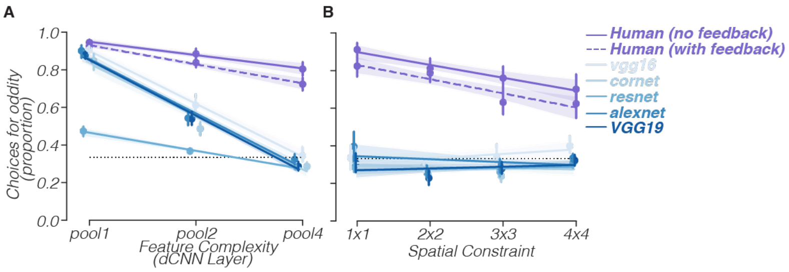


**Supp. Fig. S4.** Replication of behavioral results using dataset collected on MTurk without correct/incorrect feedback. (A) Comparison of human behavior with feedback (solid purple line), human behavior without feedback (dashed purple line), and dCNN behavior (blue lines) as a function of feature complexity. (B) Comparison of human behavior with feedback, human behavior without feedback, and dCNN behavior as a function of spatial constraint, fixing the feature complexity at the highest level (pool4).

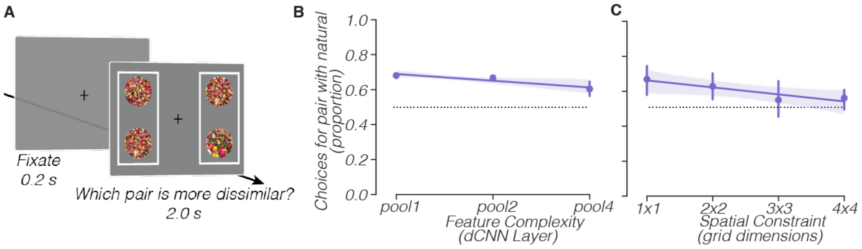


**Supp. Fig. S5**. Human behavior as a function of feature complexity and spatial constraint for the pairwise dissimilarity judgment task. (A) Task design. Subjects were shown two pairs of images and asked to select the pair which was more dissimilar from each other. (B) Human behavior as a function of feature complexity. The proportion of trials where subjects chose the pair with the natural image declined as the synths had more complex visual features. (C) Human behavior as a function of spatial constraint. The proportion of trials where subjects chose the pair with the natural declined as the arrangement of features in the synths was more strongly spatially constrained.

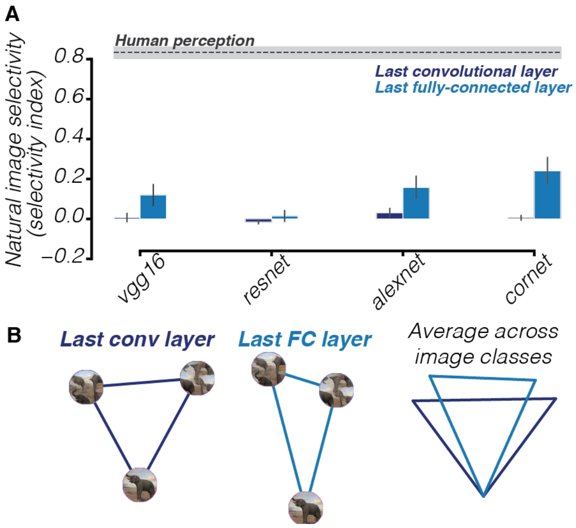


**Supp. Fig. S6**. Natural image selectivity for different dCNNs, comparing last convolutional layer to last fully-connected layer. (A) In all but one dCNN, selectivity for natural feature arrangement increases from the last convolutional layer to the last fully-connected layer. (B) Representational geometry comparing last convolutional layer to last fully connected layer.

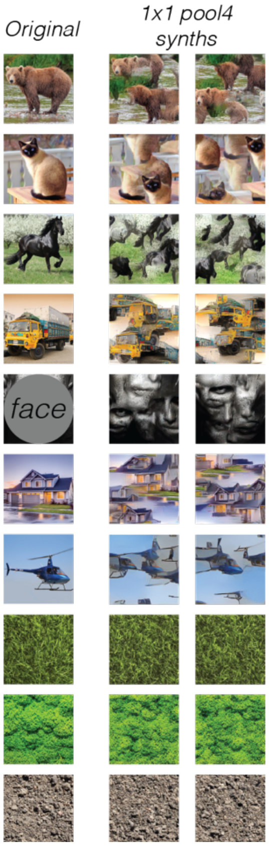


**Supp. Fig. S7**. All stimuli used in neuroimaging experiment: 10 image classes consisting of 1 natural image and 2 synths (1x1 pool4 condition) per image class.

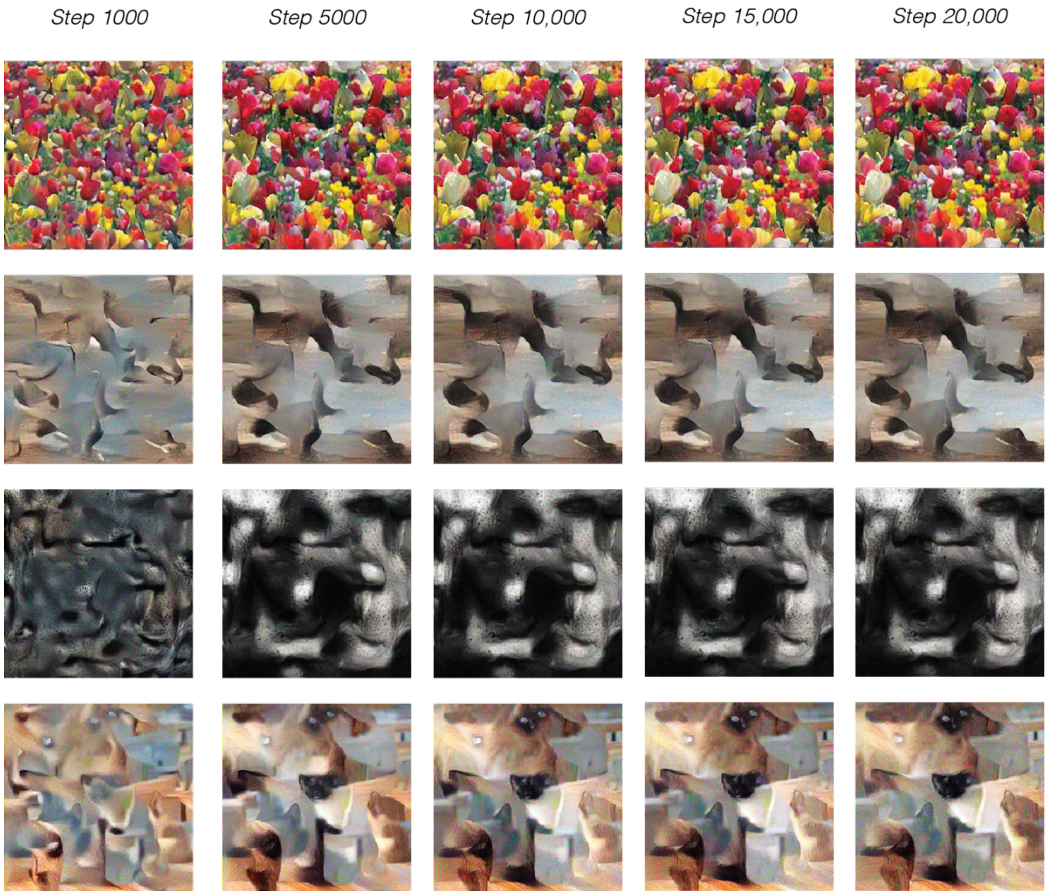


**Supp. Fig. S8**. Examples of synthesized images at different numbers of iterations in the synthesis process.

### Extended References


1. L. Gatys, A. S. Ecker, M. Bethge, Texture synthesis using convolutional neural networks. *Advances in neural information processing systems* **28**, 262–270 (2015).
2. K. Simonyan, A. Zisserman, Very Deep Convolutional Networks for Large-Scale Image Recognition. *Arxiv* (2014).
3. D. C. Liu, J. Nocedal, On the limited memory BFGS method for large scale optimization. *Math Program* **45**, 503–528 (1989).
4. J. Portilla, E. P. Simoncelli, A Parametric Texture Model Based on Joint Statistics of Complex Wavelet Coefficients. *Int J Comput Vision* **40**, 49–70 (2000).
5. J. Freeman, E. P. Simoncelli, Metamers of the ventral stream. *Nat Neurosci* **14**, 1195–201 (2011).
6. U. Güçlü, M. A. J. van Gerven, Deep Neural Networks Reveal a Gradient in the Complexity of Neural Representations across the Ventral Stream. *J Neurosci* **35**, 10005–10014 (2015).
7. C. Olah, A. Mordvintsev, L. Schubert, Feature Visualization. *Distill* **2** (2017).
8. S. A. Cadena, *et al.*, Deep convolutional models improve predictions of macaque V1 responses to natural images. *Plos Comput Biol* **15**, e1006897 (2019).
9. M. D. Zeiler, R. Fergus, Computer Vision – ECCV 2014, 13th European Conference, Zurich, Switzerland, September 6-12, 2014, Proceedings, Part I. *Lect Notes Comput Sc*, 818–833 (2014).
10. N. C. L. Kong, B. Kaneshiro, D. L. K. Yamins, A. M. Norcia, Time-resolved correspondences between deep neural network layers and EEG measurements in object processing. *Vision Res* **172**, 27–45 (2020).
11. E. P. Simoncelli, W. T. Freeman, The steerable pyramid: a flexible architecture for multi-scale derivative computation. *Proc Int Conf Image Process* **3**, 444–447 vol.3 (1995).
12. D. P. Kingma, J. Ba, Adam: A Method for Stochastic Optimization. *Arxiv* (2014).
13. M. Abadi, *et al.*, TensorFlow: Large-Scale Machine Learning on Heterogeneous Distributed Systems. *Arxiv* (2016).
14. T. S. A. Wallis, *et al.*, A parametric texture model based on deep convolutional features closely matches texture appearance for humans. *J Vision* **17**, 5 (2017).
15. P. Virtanen, *et al.*, SciPy 1.0: fundamental algorithms for scientific computing in Python. *Nat Methods* **17**, 261–272 (2020).
16. N. J. Majaj, H. Hong, E. A. Solomon, J. J. DiCarlo, Simple Learned Weighted Sums of Inferior Temporal Neuronal Firing Rates Accurately Predict Human Core Object Recognition Performance. *J Neurosci Official J Soc Neurosci* **35**, 13402–18 (2015).
17. D. L. K. Yamins, *et al.*, Performance-optimized hierarchical models predict neural responses in higher visual cortex. *P Natl Acad Sci Usa* **111**, 8619–24 (2014).
18. M. Schrimpf, *et al.*, Brain-Score: Which Artificial Neural Network for Object Recognition is most Brain-Like? *Biorxiv*, 407007 (2020).
19. D. L. K. Yamins, J. J. DiCarlo, Using goal-driven deep learning models to understand sensory cortex. *Nat Neurosci* **19**, 356–65 (2016).
20. B. A. Richards, *et al.*, A deep learning framework for neuroscience. *Nat Neurosci* **22**, 1761–1770 (2019).
21. S. Ogawa, T. M. Lee, A. R. Kay, D. W. Tank, Brain magnetic resonance imaging with contrast dependent on blood oxygenation. *Proc National Acad Sci* **87**, 9868–9872 (1990).
22. S. O. Dumoulin, B. A. Wandell, Population receptive field estimates in human visual cortex. *Neuroimage* **39**, 647–660 (2008).
23. B. A. Wandell, J. Winawer, Imaging retinotopic maps in the human brain. *Vision Res* **51**, 718–737 (2011).
24. J. L. Gardner, E. P. Merriam, J. A. Movshon, D. J. Heeger, Maps of visual space in human occipital cortex are retinotopic, not spatiotopic. *J Neurosci Official J Soc Neurosci* **28**, 3988–99 (2008).
25. A. Stigliani, K. S. Weiner, K. Grill-Spector, Temporal Processing Capacity in High-Level Visual Cortex Is Domain Specific. *J Neurosci* **35**, 12412–12424 (2015).
